# Peripherally targeted analgesia via AAV-mediated sensory neuron-specific inhibition of multiple pronociceptive sodium channels in rat

**DOI:** 10.1101/2021.10.05.463243

**Authors:** Seung Min Shin, Brandon Itson-Zoske, Chensheng Qiu, Mahmudur Rahman, Uarda Gani, Fan Fan, Theodore R. Cummins, Quinn H. Hogan, Hongwei Yu

## Abstract

This study reports that targeting intrinsically disordered regions (IDRs) of Na_V_1.7 protein facilitated discovery of sodium channel inhibitory peptide aptamers (NaviPA) for adeno-associated virus (AAV)-mediated, sensory neuron-specific analgesia. A multipronged inhibition of I_Na1.7_, I_Na1.6_, and I_Na1.3_, but not I_Na1.5_ and I_Na1.8_ was found for a prototype, named NaviPA1, which was derived from the Na_V_1.7 intracellular loop 1 and is conserved among the TTXs Na_V_ subtypes. NaviPA1 expression in primary sensory neurons (PSNs) of dorsal root ganglia (DRG) produced significant inhibition of TTXs I_Na_ but not TTXr I_Na_. DRG injection of AAV6-encoded NaviPA1 significantly attenuated evoked and spontaneous pain behaviors in both male and female rats with neuropathic pain induced by tibial nerve injury (TNI). Whole-cell current-clamp of the PSNs showed that NaviPA1 expression normalized PSN excitability in TNI rats, suggesting that NaviPA1 attenuated pain by reversal of injury-induced neuronal hypersensitivity. Immunohistochemistry revealed efficient NaviPA1 expression restricted in PSNs and their central and peripheral terminals, indicating PSN-restricted AAV biodistribution. Inhibition of sodium channels by NaviPA1 was replicated in the human iPSC-derived sensory neurons. These results summate that NaviPA1 is a promising analgesic lead that, combined with AAV-mediated PSN-specific block of multiple TTXs Na_V_s has potential as peripheral nerve-restricted analgesic therapeutics.

## Introduction

Voltage-gated sodium channels (Na_V_s) are key regulators of neuronal excitability and pain sensations (1). Mammals possess nine isoforms of Na_V_s, of which Na_V_1.7, Na_V_1.8, and Na_V_1.9 are preferentially expressed in the primary sensory neurons (PSNs) of dorsal root ganglia (DRG) (2). The prominent roles of these Na_V_ isoforms to human pain have been validated (2). Na_V_1.6, Na_V_1.1, and Na_V_1.3 are also expressed in PSNs and have been reported as possible targets for analgesics (3, 4). Currently, Na_V_1.7 is the leading target among Na_V_s for developing analgesic therapies (5).

Numerous efforts have been made over the last decades to develop selective and effective Na_V_1.7 blockers to treat pain (6), but the success has been limited. Most of the available small-molecule Na_V_1.7 blockers tested to treat pain are insufficient in target engagement, lack of targeting specificity or selective bioavailability in pain axis, and their global distribution contributes to cardio-toxicity, motor impairments, and CNS side-effects (6). Development of biologics targeting Na_V_1.7 is an alternative growing-trend (7, 8) for analgesia. Na_V_1.7 neutralizing monoclonal antibodies have analgesic efficacy, but the results are inconsistent (9). Tarantula peptide Na_V_1.7 blockers are effective analgesics but have poor membrane permeability, inadequate Na_V_1.7 selectivity, and short half-lives (6). Na_V_1.7-RNAi (6) and CRISPR-dCas9 or ZEN epigenetic Na_V_1.7 suppression for analgesic gene therapy has been proposed (10), but these interventions at mRNA and epigenetic levels lack the specificity of direct channel intervention, reducing safety and permitting off-target effects (6, 11), and anti-Cas9 immunity creates additional challenge for CRISPR gene therapies (12).

Small peptides derived from pronociceptive ion channels as functionally interfering peptide aptamers (iPA) are highly effective and selective, allowing block of specific nociceptive signaling (13, 14). Intrinsically disordered regions (IDRs) of ion channel proteins are commonly engaged in promiscuous interactomes, which are important players in multiple signaling regulations and are recognized as new and promising drug targets (15). We speculated that Na_V_1.7-IDRs contain short functional IDR domains that could play critical roles in modulating Na_V_1.7 functions and can be developed as Na_V_1.7iPAs (1.7iPA). Furthermore, the substantial conservation of Na_V_ subtype sequences implies that a given 1.7iPA could interact with other Na_V_ subtypes that have homologous sequences to Na_V_1.7 and thereby enable multipronged engagement of Na_V_ subtypes. Because multiple PSN-Na_V_s contribute to nociceptive electrogenesis and pain pathogenesis, it is conceivable that AAV-mediated expression of such multipronged NaviPA restricted in DRG-PSNs to inhibit several pronociceptive Na_V_s could be an analgesic advantage compared to block of only a single Na_V_ subtypes (16).

We here describe a novel strategy by which highly selective and nontoxic NaviPAs were designed and developed from Na_V_s-IDRs. A prototypic NaviPA1 derived from Na_V_1.7 intracellular loop 1 and conserved in TTXs Na_V_ subtypes showed multipronged inhibition of Na_V_1.7, Na_V_1.6, and Na_V_1.3 channels. NaviPA1 expression in rat PSNs rendered significant TTXs but not TTXr I_Na_ inhibition. AAV-mediated NaviPA1 expression selectively in the PSNs responsible for pain pathology in rat model produced efficient analgesia while avoiding off-site biodistribution that causes side-effects. Together, these results indicate that AAV-mediated PSN-specific, combined block of multiple nociceptive Na_V_s, has potential for future therapeutic development.

## Results

### In silico design of 1.7iPAs from Na_V_1.7-IDRs

The candidate iPAs were designed through a priori strategy aimed to define the short linear functional disordered peptides from the intrinsically disordered domains (IDDs) (13), initially from Na_V_1.7 protein IDRs, on the hypothesis that Na_V_1.7 IDDs contain the functional sequences that modulate Na_V_1.7 channel function. We analyzed the full-length of the rat Na_V_1.7 protein sequence using DisorderEd PredictIon CenTER (DEPICTER), which combines 10 popular algorithms for IDR predictions within the primary sequence, based on amino acid (aa) biophysical features for the protein’s disordered ensemble (17). Results return a score between 0 and 1 for each residue, indicating the degree to which a given residue is part of an ordered or disordered region (residues with scores >0.5 are considered as disordered). Results revealed clear order-to-disorder transitions where Na_V_1.7 transmembrane (TM) domains and intracellular portions join, and scores indicate a disordered nature of Na_V_1.7 intracellular and terminal regions (**Fig. 1A-1C**). Specifically, the most extensive IDRs are in the intracellular loops (ICL), while protein TM domains are highly ordered.

**Fig. 1.**
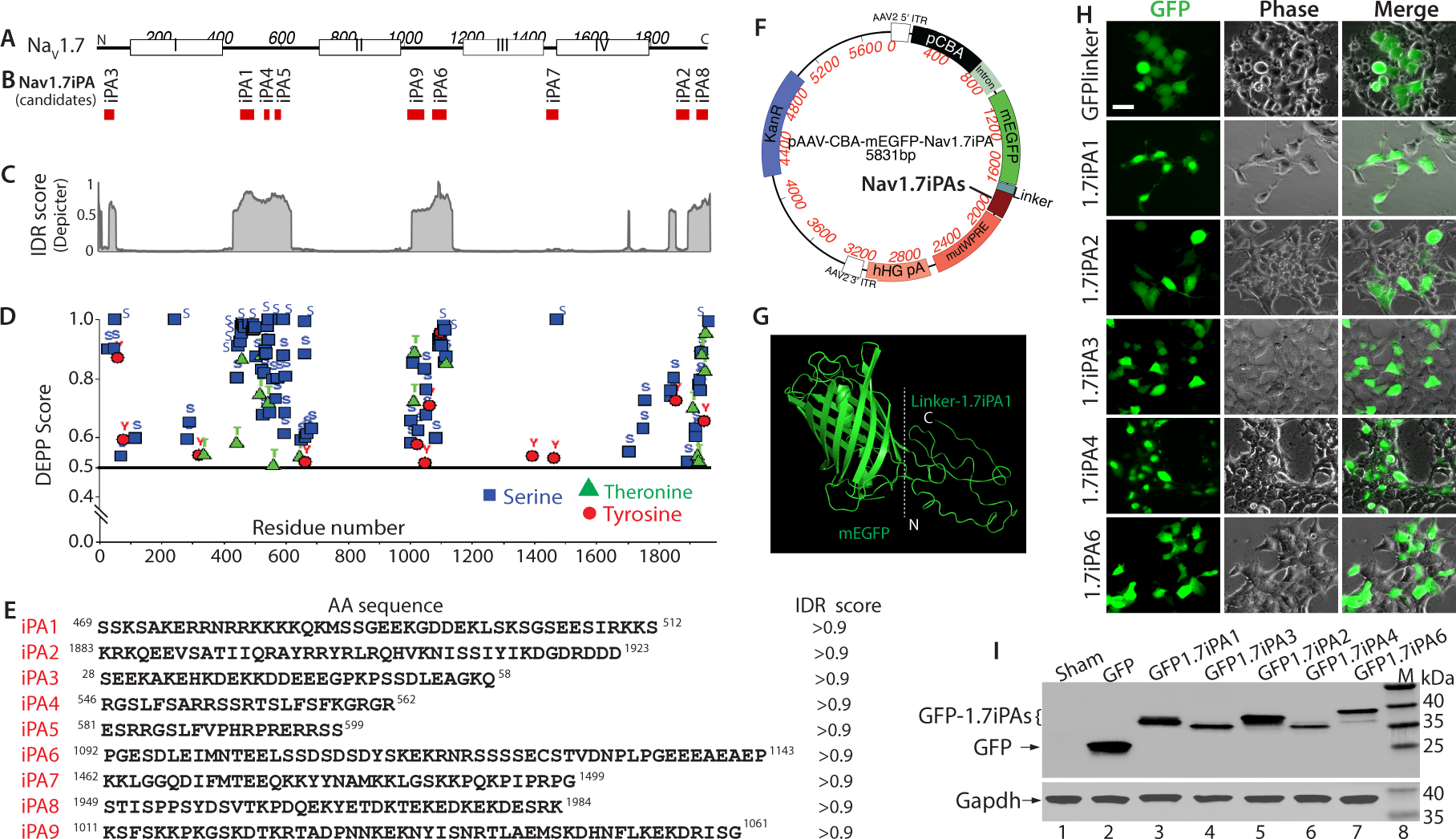
*In silico* prediction of Na_V_1.7-IDRs and design of candidate Na_V_1.7iPAs (1.7iPAs). Diagram of rat Na_V_1.7 protein, with white boxes labeling DI-DIV of Na_V_1.7 (**A**) and the red bars below showing the position of the predicted iPAs (**B**). Consensus prediction of IDRs by DEPICTER (**C**). The phosphorylation sites were predicted by DEPP (**D**). Nine candidate iPAs with their aa sequences, position in Na_V_1.7, and IDR scores (**E**). A map showing each component of an AAV plasmid coding GFP-iPA, a black line pointing to iPAs (**F**). The structure analysis of GFP1.7iPA1 by I-TASSER (**G**). Images (GFP, left; phase, middle; and merged pictures, right) show expression of constructs carrying 1.7iPA1-4 and 6 after transfection to HEK cells (**H**); Scale bar: 25μm for all. GFP and Gapdh western blots of the cell lysates after transfection with 1.7iPA1-4 and 6 to HEK cells (**I**).

Potential phosphorylation sites in the Na_V_1.7 sequence were identified using Disorder Enhanced Phosphorylation Predictor (DEPP) (18). Results showed that most potential phosphorylation residues (serine, threonine, and tyrosine with high DEPP scores) reside in Na_V_1.7-IDRs, particularly in the IDRs within the ICL1 and ICL2 (**Fig. 1D**). Na_V_1.7-IDRs feature as potential protein-protein interaction (PPI) binding sites, suggesting these IDRs could contain key binding motifs or domains of the Na_V_1.7 regulatory signaling interactome (19). These observations predict that focusing on the Na_V_1.7-IDRs could be an avenue for identifying short peptides effective in modulating Na_V_1.7 channel function.

The potentially functional domains within the Na_V_1.7-IDRs (20) were further analyzed using SLiMPrints (http://bioware.ucd.ie/slimprints.html) (21), which predict short linear motifs (SLiMs) based on strongly conserved primary aa sequences, followed by filtering based on the prediction scores (21). The enumerated motifs predicted within Na_V_1.7-IDRs suggest many possible functional peptides as ‘hot-spots’ of functional IDDs, including proteolytic cleavage sites, ligand binding sites, PTM sites, and sub-cellular targeting sites. Nine peptides were designed computationally based on IDR scores and phosphorylation sites, and were the focus as 1.7iPA candidates for further testing (**Fig. 1E, 1B**).

### Constructs of 1.7iPAs and transfection expression

AAV expression plasmids containing transgene expression cassettes encoding various GFP-1.7iPA chimeras were constructed. Specifically, the sequences for interchangeable iPA peptides were cloned with a linker sequence (GLRSRAQASNSAVDGTAGPGS) as we described previously (22), to form a chimeric transgene in a GFP-linker-iPA orientation transcribed by a hybrid human cytomegalovirus (CMV) enhancer/chicken β-actin (CBA) promoter. This generated pAAV-CBA-GFP-1.7iPAs (pAAV-1.7iPA) expression plasmids, in which the oligonucleotide encoding the interchangeable 1.7iPAs are inserted at the 3′ end of GFP (**Fig. 1F**). The predicted protein structure analysis of GFP1.7iPA1 by I-TASSER tool (https://zhanglab.ccmb.med.umich.edu/I-TASSER/) (23) shows an unfolded and extended, highly flexible structural ensemble of linker-1.7iPA1 (**Fig. 1G**), which is compatible with a well-exposed mode binding to targets. Similar structures were also identified by I-TASSER for other GFP1.7iPAs (**Suppl**. **Fig**. **1**).

### Inhibition of Na_V_1.7 current (I_Na1.7_) in HEK1.7 cells by 1.7iPAs

The stable expression of each construct was verified by transfection into HEK293 cells stably expressing human wild-type Na_V_1.7 (HEK1.7 cells), followed by immunoblots (IB). Representative tests for GFPlinker (GFP), 1.7iPAs (1, 2, 3, 4, 6) were shown (**Fig. 1H**, **I**). Initial screening experiments by whole-cell voltage-clamp of I_Na1.7_ in HEK1.7 cells, transfected with plasmids encoding nine 1.7iPAs (1.7iPA1-9), were performed to characterize the I_Na1.7_. The presence of 9 different 1.7iPAs in HEK1.7 cells on peak I_Na1.7_ density (3 days after transfection) were summarized in **Fig. 2A**, in which the data points recorded by at least 2 replicates were combined. The results showed that 1.7iPA1, 4, and 6 produced ∼68%, ∼59%, and ∼54% reduction of peak I_Na1.7_ density, respectively, while 1.7iPA2 increased peak I_Na1.7_ density (∼35%). Transfection with plasmids expressing the GFPlinker and 1.7iPA3, 5, 7, 8, and 9 showed no significant effects on peak I_Na1.7_ density, compared to sham transfected HEK1.7 cells. These experiments thus identified 1.7iPA1 and 1.7iPA4 (both derived from ICL1), as well as 1.7iPA6 (from ICL2), as effective iPAs (>50% I_Na1.7_ inhibition) (**Fig. 2B-J**). We next selected 1.7iPA1 as a prototype as it had higher inhibition of I_Na1.7_, for further ‘hit to lead’ characterization. Since 1.7iPA1 is highly conserved in aa sequences among TTXs Na_V_s and inhibits multiple TTXs I_Na_ (see further), we hereafter referred to it as NaviPA1.

**Fig. 2.**
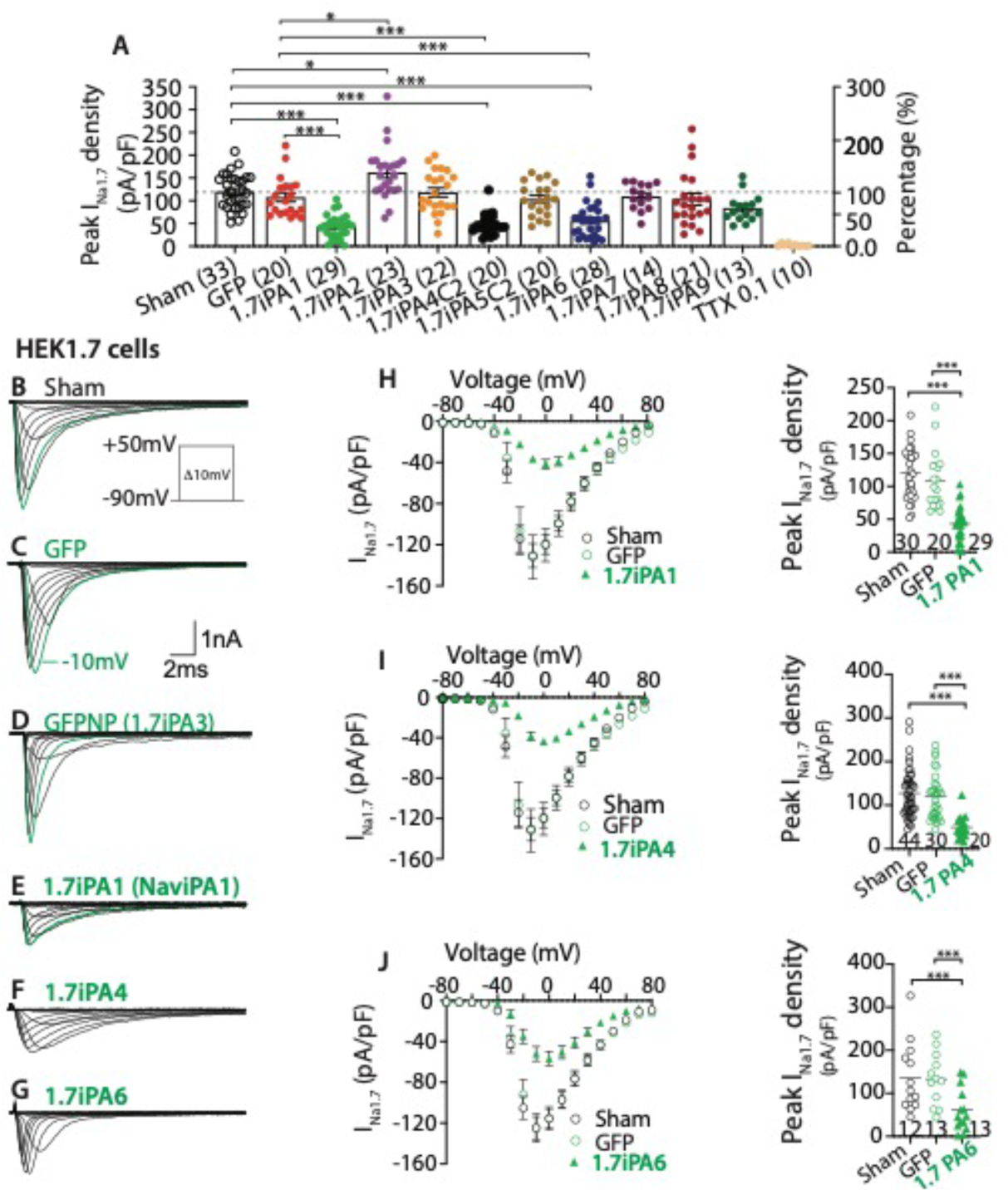
Inhibition of I_Na1.7_ of HEK1.7 cells by 1.7iPA1, 4, and 6. (**A**) Summary of the effects of 9 candidate iPAs expression in HEK1.7 cells on peak I_Na1.7_ density. (**B**-**G**) Representative traces of I_Na1.7_ recorded from sham (**B**), GFP (**C**), 1.7iPA3 (**D**), 1.7iPA1(**E**), 1.7iPA4 (**F**), and 1.7iPA6 (**G**) transfected HEK1.7 cells. Inserts: recording protocol and current/time scales. (**H**-**J**) Panels from left to right show the comparisons of I_Na_ I/V curves and peak I_Na_ density, respectively, of sham-, GFP-, 1.7iPA1(**H**)-, 1.7iPA4 (**I**)-, and 1.7iPA6 (**J**)-transfected HEK1.7 cells. *, **, *** denote p<0.05, 0.01, and 0.001, respectively, one-way ANOVA and turkey post hoc.

### Specificity of NaviPA1 occupancy to various voltage-gated ion channels

#### Development of Na_V_1.8 stable expression system based on HEK cells

To assess the potential of NaviPA1 in affecting I_Na_ conducted by Na_V_1.8 channels, we developed stable expression of recombinant human Na_V_1.8 heterologous systems based on HEK cells (HEK1.8). Stable Na_V_1.8 expression was confirmed by immunoblots of Na_V_1.8α and Na²2 in the cell lysate after at least 6 rounds of G418 selection (400-800 µg/mL), followed by single cell isolation using BIOCHIPS Single-cell Isolation Chip (ThermFisher, Rockford, IL). Both Na_V_1.8α and Na²2 are highly enriched in the plasma membrane. Functional Na_V_1.8 expression was identified by the presence of slowly inactivating inward I_Na_ elicited by voltage steps from −140 mV to +80 mV during the whole-cell voltage-clamp recordings and the averaged peak I_Na1.8_ density in ∼85% of the HEK1.8 was ∼1.0 nA, and I_Na1.8_ was sensitive to a Na_V_1.8 channel blocker, A803467 (Alomone, Jerusalem, Israel). In comparison, I_Na1.8_ amplitudes in CHO-Nav1.8 cells were generally less than 100 pA, which was insufficient for our experimental needs (**Suppl**. **Fig**. **2**).

#### Selectivity of NaviPA1 on ion channel occupancy

Na_V_ subtype stable cell lines based on HEK cells used for this experiment included HEK1.3, 1.6, 1.5, and 1.8. Sequence alignments identified substantial homology of NaviPA1 with the corresponding sequences of TTXs Na_V_1.3 and 1.6, but much less homologous to TTXr Na_V_1.5, 1.8, and 1.9 (**Fig**. **3A, B**). The NaviPA1 peptide is polyampholytic, enriched with 38.6 % positively charged arginine or lysine (17/44), and is highly conserved between rodent and human (**Fig. 3C**). Searching databases showed that two serine phosphorylation and two lysine acetylation were assigned in high throughput (proteomic discovery mass spectrometry) papers (https://www.phosphosite.org) (24), a nuclear localization signal was predicted by SeqNLS (http://mleg.cse.sc.edu/seqNLS/) (25). These analyses strongly suggest that NaviPA1 is a functional IDD peptide that has not previously been reported in the literature.

**Fig. 3.**
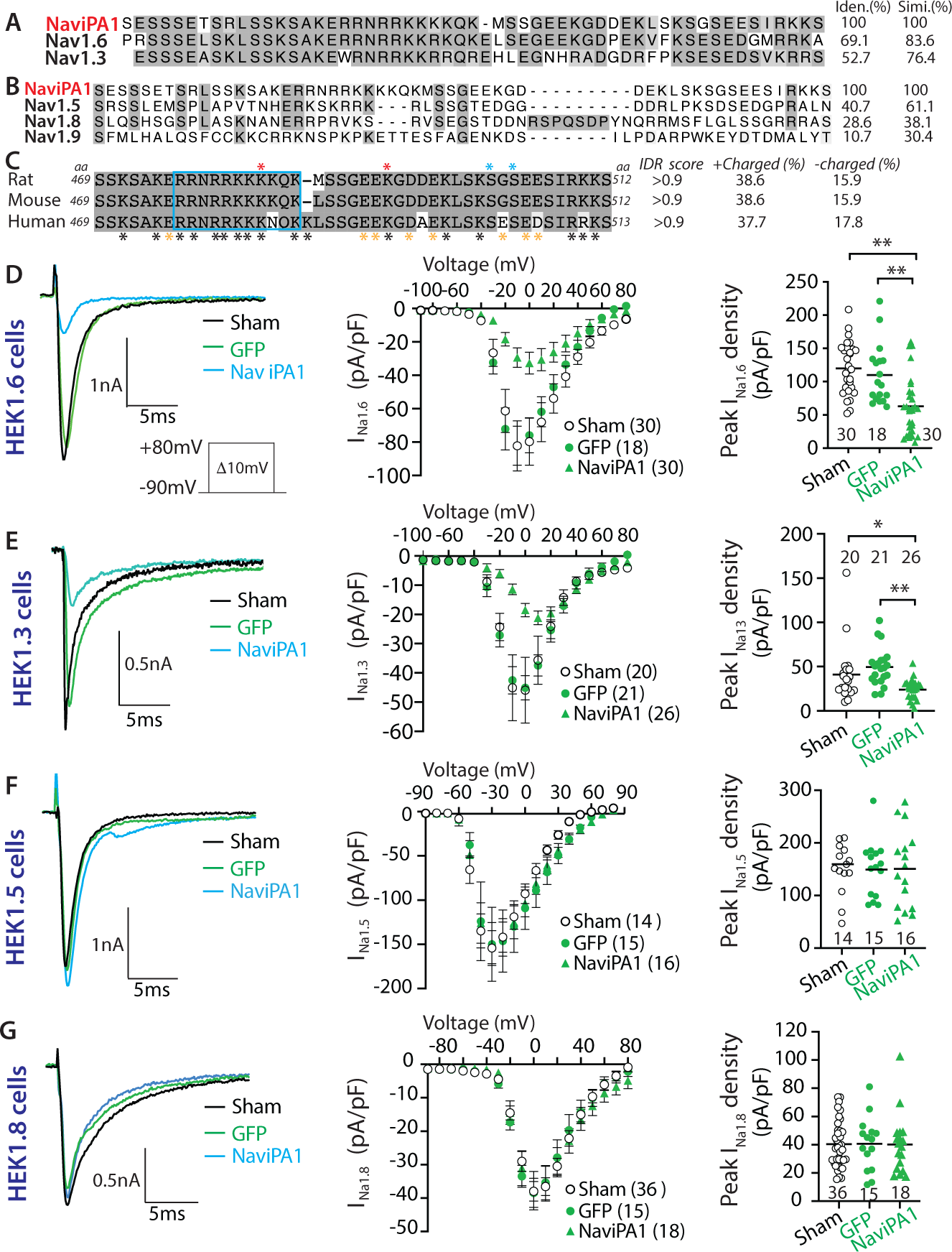
Sodium channel specificity of NaviPA1 (1.7iPA1) inhibition. Na_V_1.7iPA1 was hereafter referred to as NaviPA1 since it inhibit multiple TTXs Na_V_s. The aa sequence alignment of NaviPA1 with the corresponding sequences of TTXs Na_V_1.6 and Na_V_1.3 (**A**), as well as TTXr Na_V_1.5, Na_V_1.8, and Na_V_1.9 (**B**) of rat. The homologous aa (identity and similarity) was highlighted in heavy or light black shadows and % of identical or similar aa shown at the right sides of the alignments. NaviPA1 is highly conserved in rat, mouse, and human (**C**). Black and yellow asterisks at the bottom denote positively and negatively charged aa; the red and blue asterisks on the top denote known lysine acetylation and serine phosphorylation sites, and IDR scores and % of positively (+) and negatively (-) charged aa were shown at the right sides of the alignment. (**D**-**G**) Panels from left to right show the comparisons of representative single I_Na_ traces responded to pulse protocol (insert), I/V curves, peak I_Na_ density, respectively, of sham, 1.7NP, and NaviPA1 transfected HEK1.6 (**D**), HEK1.3 (**E**), HEK1.5 (**F**), and HEK1.8 (**G**) cells. *, **, *** denote p<0.05, 0.01, and 0.001, respectively, one-way ANOVA and turkey *post hoc*.

Expression of NaviPA1 (fused to GFP) resulted in significant block of I_Na_ conducted by fast-activating and inactivating Na_V_1.3 and 1.6 (**Fig. 3D, E)**. No effects on I_Na1.5_ and I_Na1.8_ were observed in the presence of NaviPA1 in the HEK1.5 and HEK1.8 cells (**Fig. 3F-G**). We did not test NaviPA1 against Na_V_1.1 and Na_V_1.9 channels as the expression cell lines are unavailable; however, inhibition of I_Na1.1_ by NaviPA1 is expected because of sequence homology (not shown), while no effect on I_Na1.9_ is anticipated since no sequence similarity. The effects of NaviPA1 on potassium current (I_Kv_) were tested using NG108-15 cells which naturally express potassium channels (13), and effect on high-voltage activated (HVA) I_Ca_ was recorded on AAV-mediated NaviPA1 expression in DRG-PSNs. Potent I_Na1.7_ inhibition by NaviPA1 was also confirmed in neuronal NG108-15 cells and F11 DRG-neuronal-like cells that naturally express Na_V_1.7. These experiments showed no effects of NaviPA1 on either potassium channels or HVA I_Ca_ (**Suppl**. **Fig**. **3**).

#### AAV6-mediated NaviPA1 expression in DRG-PSNs inhibits TTXs I_Na_ but not TTXr I_Na._

Because no heterologous system or cell lines can fully mimic *in vivo* conditions of sensory neurons, we further tested the functional inhibition of I_Na_ by NaviPA1 in DRG-PSNs. AAV6 vectors encoding GFP-fused NaviPA1 was generated and injected into lumbar (L) 4/5 DRG of naïve rats (male), and acutely dissociated sensory neurons from DRG were tested at 4 weeks post-injection. AAV6 encoding GFPlinker and NP (1.7iPA3) which was derived from the N-terminus of Na_V_1.7 (**Fig. 1**) and showed no impact on I_Na_ after being transfected into HEK1.7 (**Fig. 2**) were used as the controls. A voltage protocol was adopted that demonstrates successful separation of TTXr I_Na_ and TTXs I_Na_ in dissociated DRG neurons (26, 27) (**Suppl**. **Fig**. **4**). Whole-cell voltage-clamp recordings from small/medium-sized PSNs (¢.¢35mm) showed that AAV-mediated expression of NaviPA1 produced significant inhibition of total and TTXs I_Na_ whereas it produced no significant inhibition on TTXr I_Na_ (**Fig. 4A-C**).

**Fig. 4.**
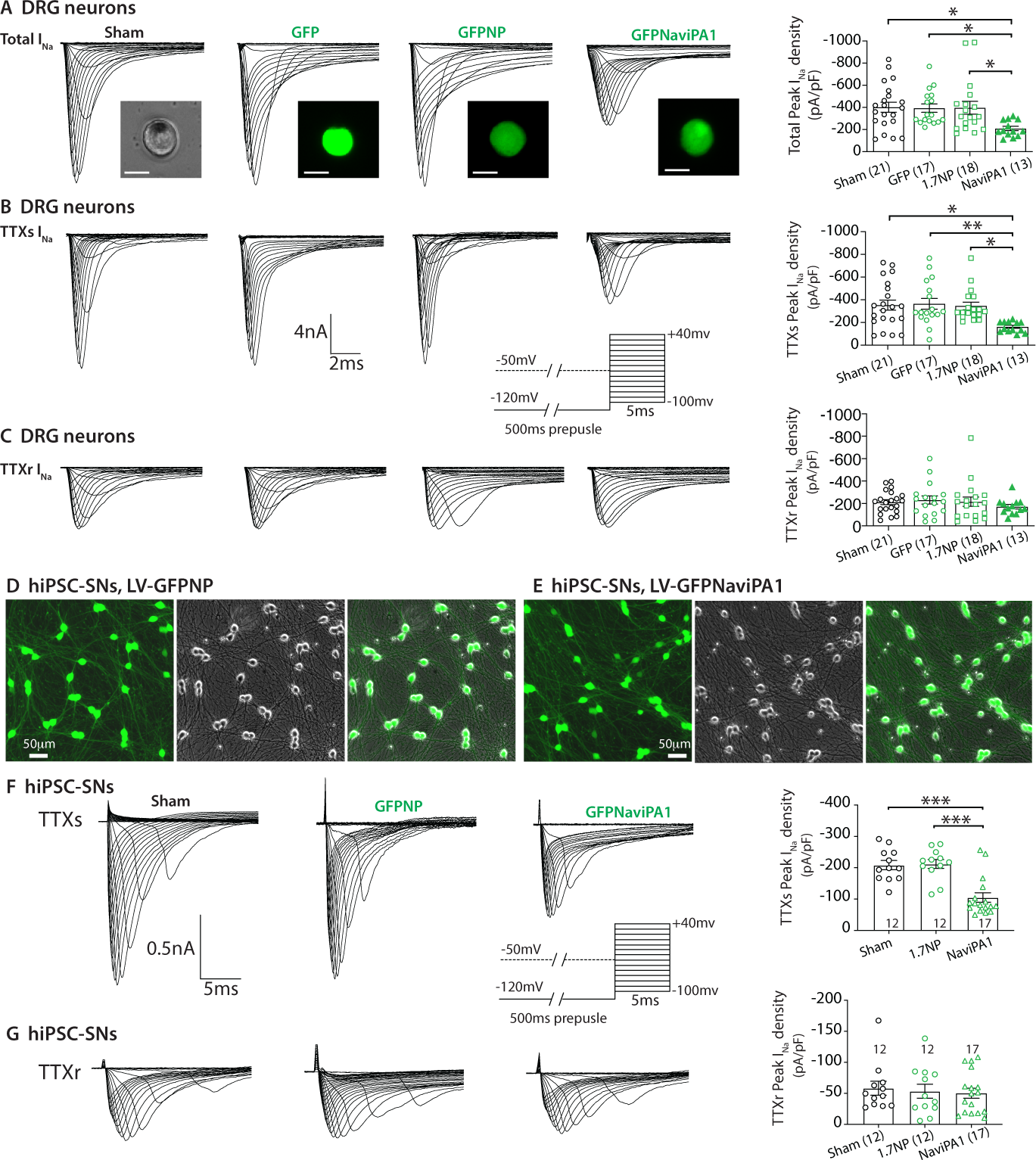
NaviPA1 on I_Na_ of rat DRG neurons (male) and hiPSC-SNs (female). (**A**-**C**) Panels from top to bottom illustrate representative traces and averaged peak I_Na_ densities of total I_Na_ (**A**), TTXs I_Na_ (**B**), and TTXr I_Na_ (**C**) recorded from sensory neurons (diameter¢.¢35mm) dissociated from naïve male rats subjected with (panels from left to right) sham, and 4wk after L4/L5 DRG injected with AAV6-encoded GFP, GFPNP, and GFPNaviPA1. Inserts: representative PSN images (scale bars 25μm for all) of each group, current/time scales, and recording pulse protocol. (**D**, **E**) Representative montage ICC images illustrate hiPSC-SNs at DIV8 after transduction with LV-GFPNP (**D**) and LV-GFPNaviPA1 (**E)** at equal MOI=10. (**F**, **G**) illustrate representative traces and averaged peak I_Na_ densities of TTXs I_Na_ (**F**), and TTXr I_Na_ (**G**) recorded from hiPSC-SNs expressing NP and NaviPA1. Inserts: current/time scales and recording pulse protocol. *, **, and *** denote p<0.05, 0.01, and p<0.001, one-way ANOVA and Turkey *post hoc*.

#### Inhibition of TTXs I_Na_ by NaviPA1 in human iPSC-derived sensory neurons

We used human induced pluripotent stem cells (iPSC)-derived sensory neurons (hiPSC-SNs, female, Anatomic, Minneapolis, MN) (28) to test whether inhibition of TTXs I_Na_ by NaviPA1 represents a meaningful and quantitative indices of the functional lead in human sensory neurons. The hiPSC-SNs were differentiated to small-sized PSN morphology with the soma diameter around 20∼25μm and developed extensive neurites after 4-7 days in vitro (DIV) differentiation cultures, indicating that these cells were efficiently committed to the neuronal lineage. We used lentivector (LV-GFP) (**Suppl**. **Fig**. **5**) to test hiPSC-SN transduction efficiency. We have succeeded in expressing NaviPA1 and 1.7NP (control) in the differentiated hiPSC-SNs by LV transduction at multiple of infection (MOI)=10 (**Fig. 4D, E**). EP recordings showed that NaviPA1 significantly inhibited TTXs I_Na_ but not TTXr I_Na_ in hiPSC-SNs (DIV12) (**Fig. 4F, G**), comparable to rat DRG-PSNs. Results indicate that inhibitory efficacy of NaviPA1 on TTXs I_Na_ defined in cell lines and rat DRG-PSNs are translatable to human PSNs.

### Molecular engagement of NaviPA1: an initial testing

We validated the specificity of Na_V_1.7 antibody by immunoblotting (IB) using the cell lysates prepared from stable cell lines expressing different Na_V_ isoforms, which showed detection of Na_V_1.7, but not Na_V_1.6 or Na_V_1.8 (**Fig. 5A**). By immunohistochemistry (IHC) on rat tissue sections, Na_V_1.7 expression was detected with high immunoreactive density in small/medium-sized PSNs using the Na_V_1.7 antibody, and Na_V_1.7 was also detected in spinal cord dorsal horn (SDH), sciatic nerve nodes of Ranvier (29), and cutaneous terminals in hindpaw (**Fig. 5B-E**), with the patterns similar to the prior reports (30). These results confirmed the specificity of the Na_V_1.7 antibody to detect Na_V_1.7 expression by IHC and immunoblot.

**Fig. 5.**
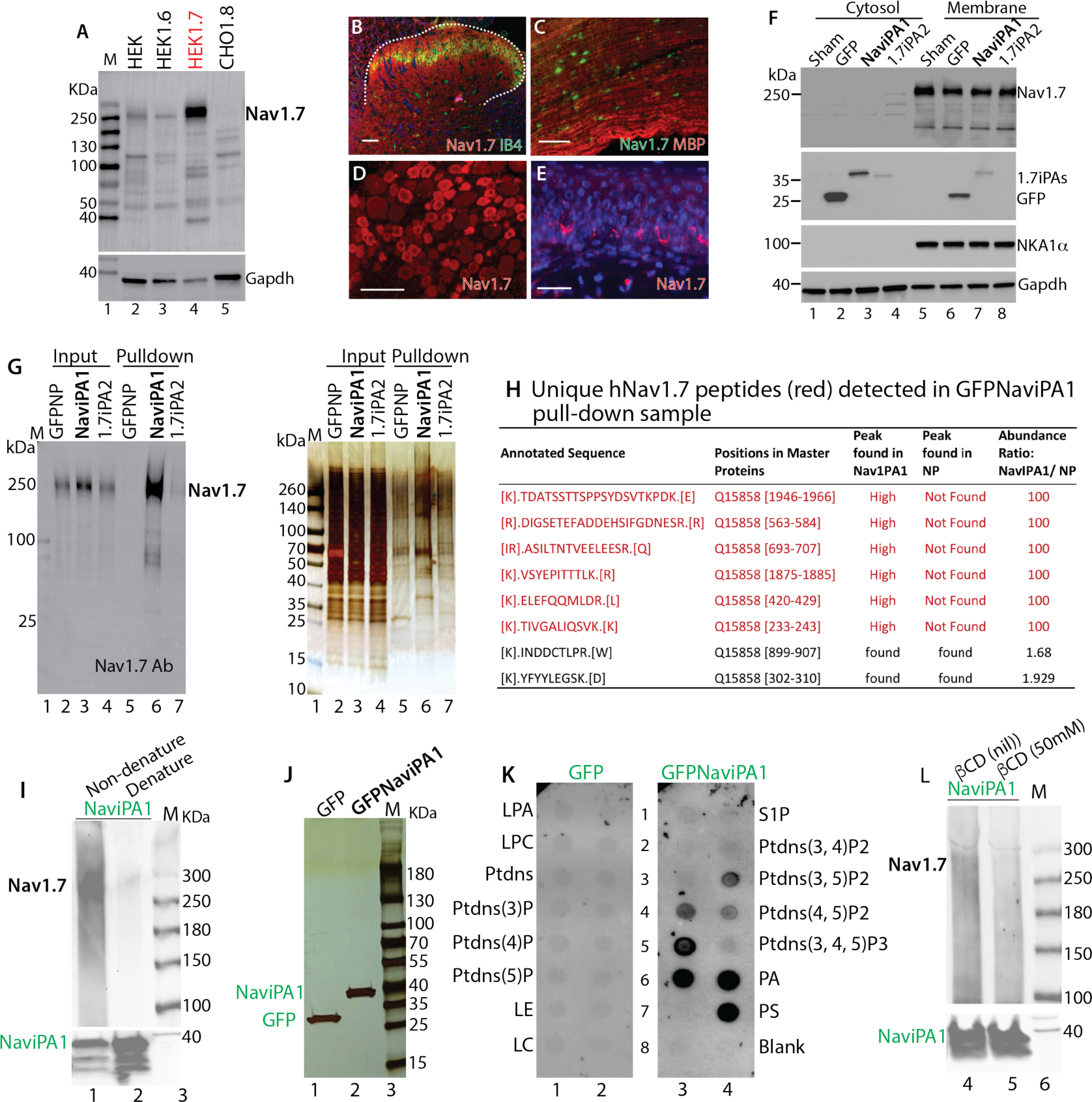
NaviPA1 binds to full-length Na_V_1.7 protein and phosphoinositides. (**A**) Immunoblots (IB) show selectivity of Na_V_1.7 antibody using cell lysates from HEK cells, and HEK1.6, HEK1.7, and CHO1.8 cells. (**B**-**E**) Representative IHC images show Na_V_1.7 detection (red) in SDH (red), sciatic nerve (green), DRG neurons (red), and cutaneous nerve fibers (red). Scale bars: 100μm. (**F**) IBs of Na_V_1.7, GFP, NKA, NKA1α, Gapdh in the cytosol and membrane samples extracted from HEK1.7 cells transfected with sham, GFP, GFPNaviPA1, and GFP1.7iPA2. (**G**) Na_V_1.7 IB (left) and silver stain (right) of inputs (cell lysates, 20μg for each lane) and pulldown beads (10μL for each lane) prepared by a ‘nondenaturing’ lysis buffer from HEK1.7 cells transfected with GFP, GFPNaviPA1, and GFP1.7iPA2. (**H**) Stained gel pieces ranging 100-300kDa (G, red asterisk denotes Na_V_1.7 site) from GFPNP and GFPNaviPA1 excised for mass spectrometry showing detection of unique human Na_V_1.7 peptides (red) in GFPNaviPA1 pull-down sample. NaviPA1-Nav1.7 interaction in non-denature and denature preparation (**I**). Silver stain on 1D SDS-PAGE gel of GFP-affinity pulldown beads in the NG108-15 cells transfected with GFPNaviPA1 and GFP and cell lysates prepared using denaturing RIPA buffer (**J**) and the results of PIP strip analysis (**K**). LPA, Lysophosphatidic acid; LPC, Lysophosphocholine; PtdIns, Phosphatidylinositol; PtdIns(3)P, PtdIns (3) phosphate; PtdIns(4)P, PtdIns (4) phosphate; PtdIns(5)P, PtdIns (5) phosphate; PE Phosphatidylethanolamine; PC, Phosphatidylcholine; S1P, Sphingosine 1-Phosphate; PtdIns(3,4)P_2_, PtdIns (3,4) bisphosphate; PtdIns(3,5)P_2_, PtdIns (3,5) bisphosphate; PtdIns(4,5)P_2_, PtdIns (4,5) bisphosphate; PtdIns(3,4,5)P_3_, PtdIns (3,4,5) trisphosphate; PA, Phosphatidic acid; PS, Phosphatidylserine. NaviPA1-Nav1.7 interaction is broken after removing lipids by ²-cyclodextrin (50mM) (**L**).

Since Na_V_1.7 is an integral membrane protein, we therefore tested whether NaviPA1 expression in the HEK1.7 cells would interrupt Na_V_1.7 intracellular trafficking. Our results do not support this mechanism since no clear reduction of membrane Na_V_1.7 protein was evident in the fractionized preparations, transfected with NaviPA1 and controls (**Fig. 5F**). Studies have shown that IDRs in the membrane proteins engage in interactions with the membrane (31). To test whether NaviPA1 interference of Na_V_1.7 might be via direct block of Na_V_1.7, GFP affinity pull-down by ChromoTek GFP-Trap (ChromoTek, Rosemont, IL) was performed after transfection of GFP-NaviPA1 in HEK1.7 cells using GFPlinker (GFP) and GFP-1.7iPA2 (**Fig. 1**) as the controls. Cell lysates were prepared by a lysis buffer containing 0.5% Nonidet p40, a ‘non-denaturing’ mild lysis detergent, for preventing interaction breaking and maximizing the retention of NaviPA1-protein interactions (32). Immunoblots verified full-length Na_V_1.7 protein trapped in the GFPNaviPA1 pull-down sample but not in controls (**Fig. 5G)**, and nLC-MS/MS detection of unique hNa_V_1.7 peptides (**Fig. 5H)** confirmed hNa_V_1.7 on the excised band from silver-stained SDS-PAGE gel of GFPNaviPA1 affinity pull-down sample. NaviPA1 binding to full-length Na_V_1.7 was broken when cell lysate prepared by use RIPA buffer containing strong detergents (see below) (**Fig. 5I)**. These results indicate that NaviPA1 block of Na_V_1.7 channel activation could be via binding to the Na_V_1.7 protein, i.e., an intra-molecular domain-domain interaction (intraDDI) (33). It has been reported that polybasic IDRs in transmembrane proteins preferably bind to negatively charged lipids (34, 35). We reasoned that NaviPA1 might be able to bind phosphoinositides, and this hypothesis was tested by using phosphatidylinositol phosphate (PIP) strips (Echelon PIP Strip, Salt Lake City, UT). GFPNaviPA1 and GFP (control) were transfected into neuronal NG108-15 cells, and cell lysates were prepared by a RIPA buffer containing 0.1% SDS and 1% Triton X100 (strong detergents) and 1% deoxycholate (anionic detergent) for maximally denaturing to break NaviPA1 PPI complex formations. Silver stain after SDS-PAGE gel showed clean purification of GFP and GFPNaviPA1 (**Fig. 5J)** and samples were applied to the PIP strips. Results (**Fig. 5K**) showed that GFP NaviPA1 were efficiently bound to a number of anionic PIPs, PIP2, phosphatidic acid (PA), and phosphatidylserine (PS). In contrast, affinity pull-down GFP did not show clear binding to lipid spots as previously reported (36). NaviPA1 binding to Na_V_1.7 was broken after removing lipids by exposure GFP-pull down sample to ²-Cyclodextrin (²CD) (repeated two times), suggesting a possibility that polybasic NaviPA1 peptide functions via a pathway interacting with membrane lipid networks correlated with the Na_V_1.7 protein function. This is supported by reports that basic residues, often clustered in IDRs, can modulate membrane protein functions by binding via electrostatic interactions with lipids (37, 38).

Delineation of the molecular mechanisms underlying these events in future studies is of interests from both pathophysiological and therapeutic perspectives. Our goal of this study is to develop a strategy of peripherally-targeted analgesia via AAV-mediated sensory neuron-specific inhibition of sodium channels. Therefore, in the following in vivo experiments, we focused on testing whether DRG-PSN-targeted expression of NaviPA1 is effective in attenuating neuropathic pain behaviors.

### Analgesia after intraganglionic delivery of AAV-NaviPA1 in rats after TNI

We first conducted a pilot in vivo analgesia testing. High-titer and high-purity of AAV6-GFPNaviPA1 (AAV6-NaviPA1) and control AAV6-GFPNP (AAV6-NP) were generated and injected into the L4/5 DRG of adult male rats. Three weeks after DRG-AAV injection, TNI was induced, and subsequent sensory behavior evaluation was performed on a weekly basis for an additional 5 weeks, after which tissues were harvested for IHC characterization of transgene expression. Results (**Suppl. Fig**. **6**) showed that AAV6-NaviPA1 injection reduced TNI-induced mechanical and cold sensitization. IHC revealed efficient NaviPA1 (fused to GFP) expression in DRG neurons and their peripheral (cutaneous) and central terminals (SDH). These data indicate that sustained expression of the NaviPA1 selectively in the PSNs of the pathological DRG after TNI prevented development of pain behaviors.

### Treatment of established neuropathic pain by DRG-AAV6-NaviPA1 in male rats

We next extended experiments to evaluate the effectiveness of DRG-AAV6-NaviPA1 in a more clinically relevant design for reversal of established pain behaviors, including both evoked responses as well as spontaneous ongoing pain following TNI. In the experimental design, the sensitivity to mechanical and thermal cutaneous stimulation was assessed at baseline and weekly after TNI for 2 weeks before AAV injection. Thereafter, rats were randomized to receive DRG injection of either AAV6-NaviPA1 or control AAV6-NP into the L4/L5 DRG ipsilateral to TNI, after which sensory behaviors were evaluated weekly for additional 6 weeks. As a terminal experiment, Gabapentin (GBP,100mg/kg, i.p.)-induced conditioned place preference (CPP) test was performed in both groups to evaluate spontaneous pain (13, 39). Behaviors measured before AAV injection at the 14^th^ day after TNI were used as a treatment baseline (tBL) to evaluate effectiveness of vector treatments (**Fig. 6A, B)**. Tissues were harvested for IHC characterization of transgene and target gene expression, and for whole-cell current-clamp of neuronal excitability on dissociated DRG neurons.

**Fig. 6.**
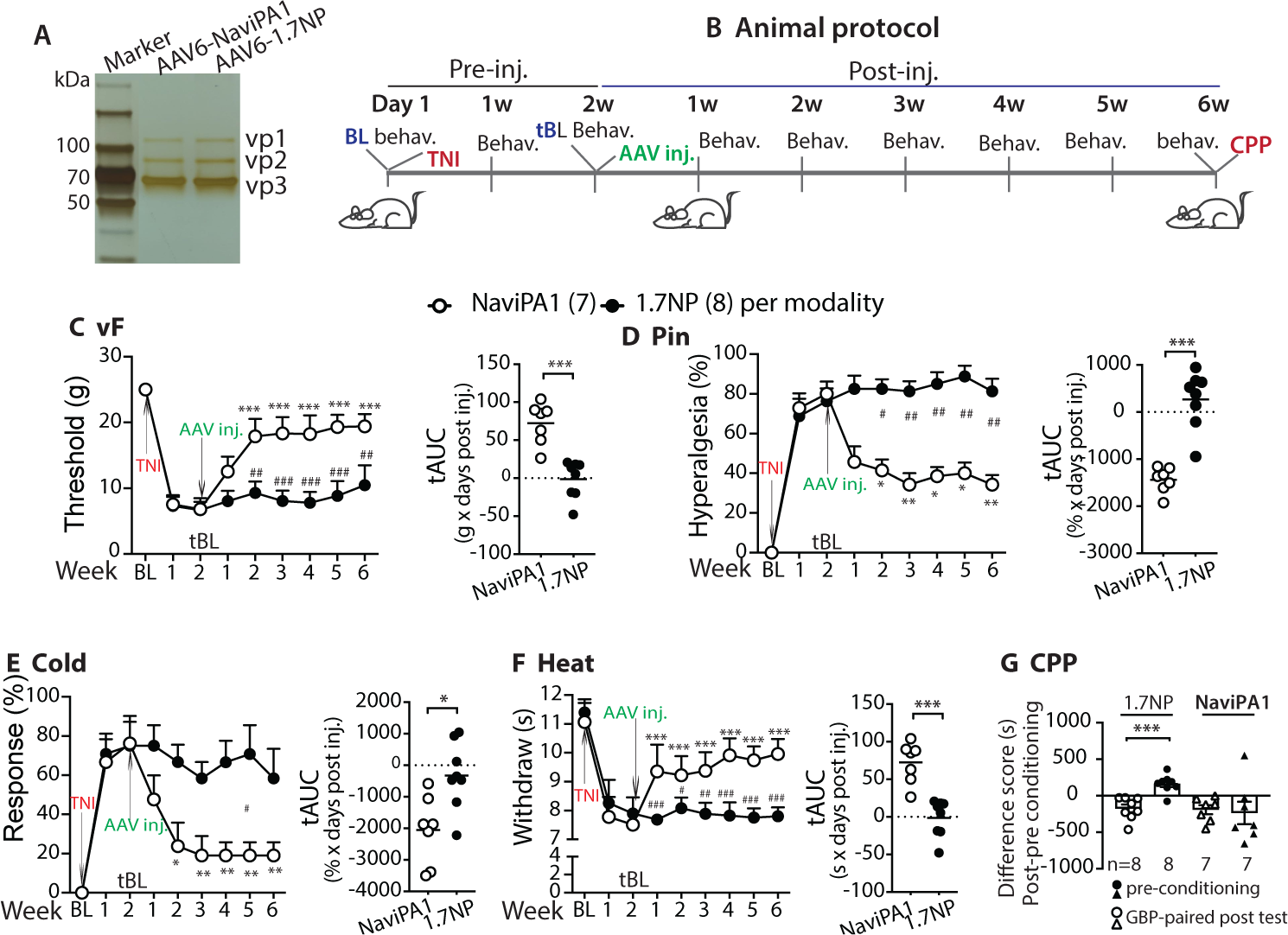
Treatment of established neuropathic pain by DRG AAV6-NaviPA1 (male rats). Purified AAVs (**A**, silver stain) were prepared for the experiment in an animal protocol schematically outlined (**B**). The time courses for the group averages of vF, Pin, Heat, and Cold before and after DRG injection of either AAV6-NaviPA1 (n=7) or AAV6-NP (control, n=8) (**C-F**); \**p*<0.05, \*\**p*<0.0,1 and \*\*\**p*<0.001 for comparisons to the tBL within group and ^#^*p*<0.05, _##_*p*<0.01, and ^###^*p*<0.001 between groups. Repeated measures two-way ANOVA for vF and Heat, and Tukey (within group) and Bonferroni (between group) *post hoc*; and non-parametric Friedman ANOVA for Pin and Cold tests and Dunn’s *post hoc*. Right panels of C to F show tAUC; \**p*<0.05 and \*\*\**p*<0.001, comparisons of tAUC between groups (unpaired, two-tailed student**’**s *t* tests). (**G)** Results of CPP scores (seconds, s) of pre-conditioning chamber and of the GBP-paired chamber between AAV-NaviPA1 (n=7) and AAV-NP (control, n=8), \*\*\**p*<0.001 (unpaired, two-tailed Student**’**s *t* test).

All rats developed multiple modalities of pain behaviors 2 weeks after TNI, including lowered threshold for withdrawal from mild mechanical stimuli (vF), more frequent hyperalgesic-type responses after noxious mechanical stimulation (Pin), and hypersensitivity to heat and acetone stimulation. These behaviors persisted after injection of the control AAV6-NP during the 6 weeks of observation course. In contrast, rats injected with AAV6-NaviPA1 showed reversal of these changes, which was maintained throughout the observation period (**Fig. 6C-F**). Using a biased CPP paradigm (40), the effect of AAV-NaviPA1 treatment on spontaneous pain was evaluated. None of the animals in either group was excluded from study because of their baseline preference/avoidance for a chamber (40). A significant GBP-induced CPP effect was observed in the TNI rats injected with AAV6-NP, while there was no significant difference in the time spent in the initially nonpreferred chamber during baseline *vs*. testing period in AAV-NaviPA1 treated TNI animals, indicating that AAV-NaviPA1 treatment significantly relieved on-going spontaneous pain (**Fig. 6G)**.

Histological examination (**Fig. 7A-G**) determined the *in vivo* transduction rate for AAV6-NaviPA1 at the 6^th^ week after vector injection. The NaviPA1-positive neurons (GFP) comprised 38 ± 3% (1283 out of 3447 total neuronal profiles) identified by a pan-neuronal marker β3-tubulin (n **=** 6 DRG, 3-4 sections per DRG, selected as every fifth section from the consecutive serial sections). Transduced DRG neurons included the full-size range of the PSNs that also expressed Na_V_1.7 and Na_V_1.6, and expression showed multiple subcellular localizations, preferably in PSN cytosol. Positive GFP signals were not detected in GFAP-positive perineuronal glial cells. GFP signals were also detected in the ipsilateral dorsal horn, sciatic nerve, and cutaneous afferent terminals.

**Fig. 7.**
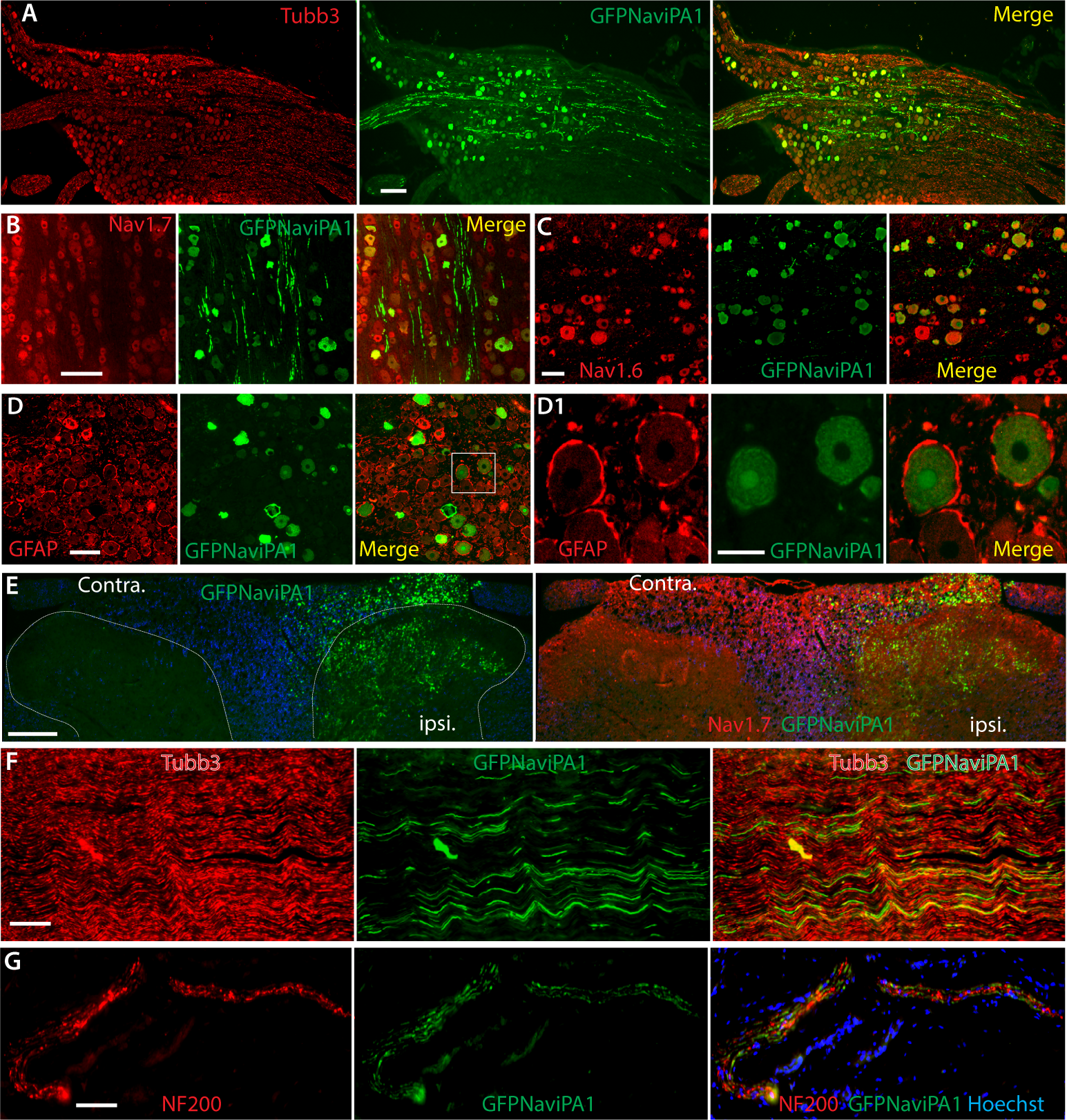
IHC of GFP-NaviPA1 and target gene expression. (**A-D**) Representative IHC montage images (GFPNaviPA1 with Tubb3) show neuronal expression profile 6 weeks after AAV-NaviPA1 injection in TNI rats (**A**), colocalization of GFP-NaviPA1 with Na_V_1.7 and Na_V_1.6 positive neurons (**B, C**), but not with GFAP positive perineuronal glia (**D**, the square region was enlarged and montage images shown as **D1**). (**E-G**) Representative IHC montage images illustrate GFPNaviPA1 (green) and Na_V_1.7 (red) in PSN central terminals of ipsilateral spinal dorsal horn (**E**), GFPNaviPA1 (green) and Tubb3 (red) in sciatic nerve (**F**), and GFPNaviPA1 (green) and NF200 (red) in PSN peripheral terminals of skin section (**G**). Scale bar (μm): A, 200; B, C, D and D1, 100; E, 200; G and G, 50μm.

These findings together demonstrate that DRG injection of AAV6-encoded NaviPA1 induced NaviPA1 expression restricted to the PSNs of injected DRG and their peripheral and central processes. This strategy via AAV6-mediated expression of NaviPA1 selective in the sensory neurons of the anatomically segmental DRG responsible for pain pathophysiology has clear analgesic effectiveness in normalizing the established peripheral hypersensitivity for both evoked and spontaneous pain behavior in the rat model of peripheral injury-induced neuropathy.

### Reversal of PSN hyperexcitability by AAV6-NaviPA1 treatment (male rats)

Increased excitability of nociceptive PSNs is a fundamental process underlying neuropathic pain (41). We therefore examined whether AAV6-NaviPA1 treatment reverses the enhanced neuronal excitability of nociceptive PSNs following TNI (42), using the whole-cell current-clamp AP recording of DRG dissociated neurons from rats after the treatment protocol shown in **Fig. 6B**. Although TNI results in DRG containing co-mingled injured and uninjured axons, nerve-injury can induce an increase of voltage-gated ion channel activity in both axotomized neurons and adjacent intact neurons, leading to similar electrophysiological (EP) changes and increased discharge frequency in axotomized and neighboring intact DRG neurons (43, 44), possibly through interneuronal signaling and coupling (45). We therefore recorded from randomly chosen small/medium-sized neurons (<35 μm in diameter) (46) in the cultures from dissociated L4 and L5 DRG. Transduced neurons were identified by GFP fluorescence, and excitability was evaluated by measuring rheobase and repetitive action potential (AP) firing during 250ms current pulses stepping from 100pA and 280pA current injection. Results showed that the averaged rheobase in the neurons from TNI rats was significantly decreased and, in response to a step stimulus, the frequency of APs evoked in neurons from TNI rats was significantly increased, compared to sham controls. These were normalized in the transduced neurons after AAV6-NaviPA1 treatment, whereas NP-transduced neurons had no significant effects (**Fig. 8)**. These findings indicate that reversal of nerve injury-induced sensory neuronal hyperexcitability by NaviPA1 may contribute to its analgesic effects in attenuation of neuropathic pain behaviors, i.e., conduction block of TTXs Na_V_ ion channels selectively in PSNs leads to a substantial decrease in neural excitability, resulting in mitigation of pain behaviors.

**Fig. 8.**
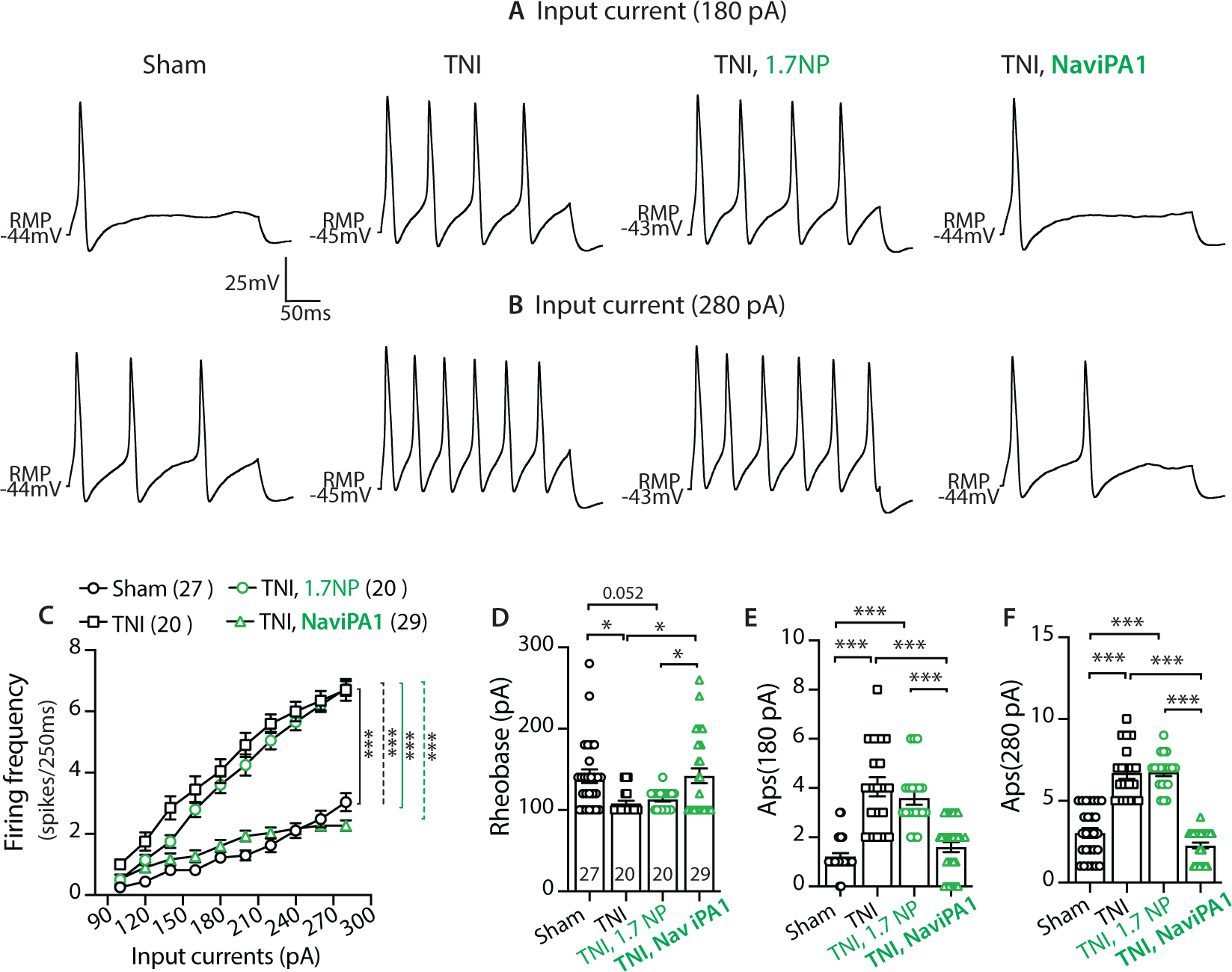
NaviPA1 expression on neuronal excitability of rat PSNs (male). (**A**, **B**) Representative AP traces elicited by 250 ms depolarizing current of 180 pA (**A**) and 280 pA (**B**) (same cells) from RMP were recorded from DRG neurons dissociated from the rats of sham, TNI only, and GFP-expressing neurons in TNI treated with AAV6-NP or AAV6-NaviPA1, as indicated. (**C**) Comparison of responses (number of APs evoked by a 250 ms stimulus) for the populations of DRG neurons in different groups across a range of step current injections from 100 to 280 pA; \*\*\**p* < 0.001, two-way ANOVA of main effects of groups with Bonferroni *post-hoc*. Scatter plots with bars show analysis of the rheobases (**D**) and AP numbers evoked by input current at 180 pA (**E**) and 280 pA (**F**) from RMP, respectively. The number in each group is the number of analyzed neurons per group. *, and ***denote *p*<0.05 and <0.001, respectively, One-way ANOVA and Turkey *post-hoc*.

### Analgesia of DRG-AAV6-NaviPA1 treatment in female TNI rats

Sex differences exist in experimental and clinical pain and in responsivity to interventions (47). We therefore next tested whether DRG-AAV6-NaviPA1 treatment is also effective in attenuating hypersensitivity induced by TNI in female animals, using the protocol similar to the tests in male animals (**Fig. 6**). The same batch preparation of AAV6-NaviPA1 and AAV6-NP tested in male rats was used for injection. Results showed that the female rats displayed similar phenotypic development of hypersensitivity after induction of TNI to male rats and that both evoked mechanical/thermal hypersensitivity and GBP-CPP responses were normalized after AAV6-NaviPA1 treatment, demonstrating comparable analgesic effects (**Fig. 9A-E**) to the male animals. IHC on the DRG sections from female TNI rats 6 weeks after AAV6-NaviPA1 injection also revealed GFP-NaviPA1 expression profile comparable to male rats (**Fig. 9F-G**), however, transduction rate in DRG neurons was not quantified. Thus, although not rigorously compared, treatment effects were comparably concordant between the sexes, suggesting that a sexual dimorphism seems not apparent for both pain behavior phenotypes after TNI and in responsivity to DRG-NaviPA1 treatment in our studies (13).

**Fig. 9.**
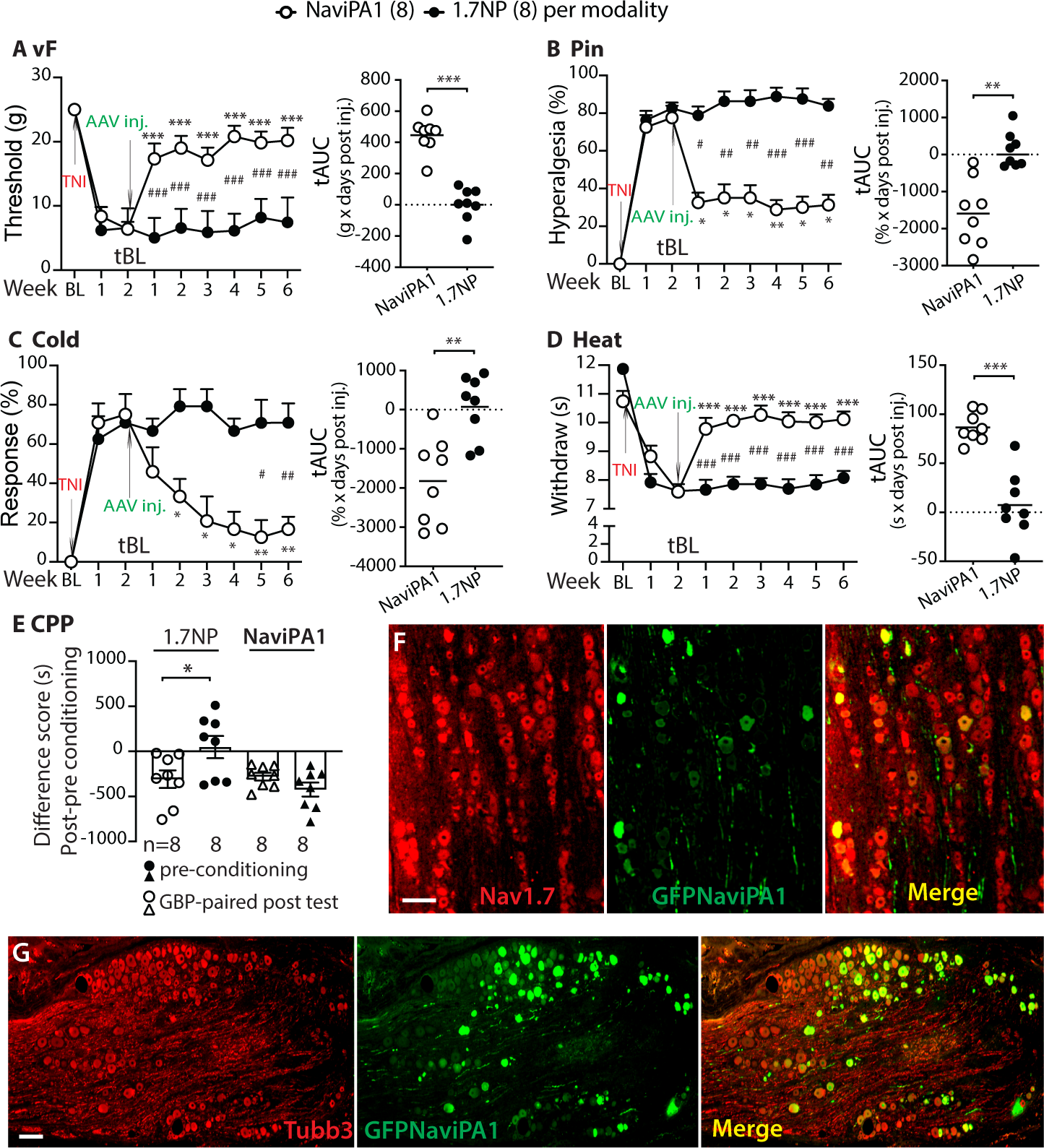
Analgesia of DRG-AAV6-NaviPA1 treatment in female TNI rats. Analogous figures to Fig. 6 show significant analgesia after DRG delivery of AAV6-NaviPA1 in the established TNI pain behaviors of female rats. *, **, and *** denote *p*<0.05, 0.01, and 0.001 for comparisons to the treatment baseline (tBL) within group and ^#^*p*<0.05, ^##^*p*<0.01, and ^###^*p*<0.001 for comparisons between groups (**A**-**D**). Repeated measures parametric two-way ANOVA for vF and Heat followed by Tukey (within group) and Bonferroni (between group) *post hoc*; and non-parametric Friedman ANOVA for Pin and Cold tests and Dunn’s *post hoc*. Right panels of **A** to **D** show tAUC calculated using measures 14-day post TNI and before vector injection as tBL; ** *p*<0.01 and *** *p*<0.001, comparisons of tAUC between groups (unpaired, two-tailed Student**’**s *t* tests). CPP difference scores (s) of pre-conditioning chamber and of the GBP-paired chamber between AAV-NaviPA1 (n=8) and AAV-NP (control, n=8), \**p*<0.01 (unpaired, two-tailed Student**’**s *t* test) (**E**). Representative montage IHC images colocalization of GFP-NaviPA1 with Na_V_1.7 (**F**) and with Tubb3 (**G**) show neuronal expression profile 6 weeks after AAV-NaviPA1 injection (**G**). Scale bar: 50μm for F and 100μm for G.

## Discussion

Sustained peripherally targeted analgesia without risk of addiction is a global unmet medical need (48). Na_V_1.7 is currently a leading target for analgesic pharmaceutics. However, ample evidence demonstrates that multiple sensory neuronal Na_V_s contribute to nociceptive electrogenesis and pain pathogenesis (16, 49). Here, we reported that targeting Na_V_-IDRs facilitated discovery of NaviPAs. A prototypic NaviPA1, initially derived from Na_V_1.7, is highly conserved in sequences among TTXs Na_V_s, and accordingly, demonstrated multipronged inhibitory feature to TTXs I_Na_ conducted by Na_V_1.7, Na_V_1.6, Na_V_1.3 but no effect on TTXr I_Na_ conducted by Na_V_1.8 and Na_V_1.5. NaviPA1 expression in DRG-PSNs produced selective inhibition of TTXs I_Na_ but not TTXr I_Na_. DRG delivery of AAV6-encoded NaviPA1 significantly attenuated established nerve injury-induced pain behaviors in male and female animals for both evoked mechanical and thermal hypersensitivity and on-going or spontaneous pain behaviors, the symptoms commonly found in patients suffering from multiple types of painful neuropathy (50). Additionally, blockade effects of TTXs I_Na_ by NaviPA1 were replicated in the hiPSC-SNs, supporting a translational potential. Because several different types of Na_V_s in sensory neurons combine to trigger nociceptor electrogenesis required for AP trains (1), block of several of these specific in DRG-PSNs is conceived to be a therapeutical advantage for neuropathic pain.

Chronic pain in almost all cases is maintained by ongoing afferent hyperactivity originating from peripheral pathological sources (51–53). Thus, development of novel peripheral acting strategies for pronociceptive Na_V_ inhibition in the PSNs would be an ideal approach for clinical pain treatment (2, 54). Our strategy described here includes a novel approach by which highly selective and nontoxic NaviPA1 is designed and developed from Na_V_-IDRs, which is delivered by using AAV to the pathological DRG. PSN-restricted inhibition of multiple pronociceptive TTXs Na_VS_ is predicted to have advantages for DRG-targeted analgesia, as a recent expert commentary states that “disappointing analgesic pharmaceutics after a single Na_V_1.7 inhibition might correlate to the facts that the excitability of neurons is determined by several different Na_V_ channels and targeting just one may not be sufficient by itself” (55). It is known that human subjects and animal models that are heterozygous for null mutations of Na_V_1.7 are normal in sensory phenotypes. Thus, AAV-mediated NaviPA1 expression restricted in DRG-PSNs may induce analgesia via a combined partial inhibition of Na_V_1.7, Na_V_1.6, and Na_V_1.3 (likely including Na_V_1.1), while avoiding undesirable side-site effects otherwise due to global distribution of small molecule inhibitors. A complete block of Na_V_1.7 activity is not intended since it may induce a state of total insensitivity to pain where unintended self-injury would occur (56).

Pain-sensing PSNs can become hyperexcitable in response to peripheral nerve injury, which in turn leads to the development of neuropathic pain. Multiple lines of evidence from both preclinical and clinical studies demonstrate that block of peripheral nociceptive input can effectively relieve pain symptoms including spontaneous pain (57, 58). Therefore, treatments targeting the peripheral PSNs both avoid CNS side effects and also are likely to succeed. Indeed, a recent expert commentary states that “activity in primary afferent neurons represents a ‘low-hanging target’ in the development of safe therapies” for patients with chronic pain (51). Delivering drugs to the DRG is well developed and safe, for instance as used by anesthesiologists for regional blockade and by pain physicians for diagnosis and treatment of radiculopathy (59). Injection into the DRG has minimal consequences in preclinical models (60). It has also been demonstrated that unintentional intraganglionic injection commonly accompanies clinical transforaminal epidural steroid injection (59), a very common procedure with minimal risk of nerve damage. Thus, the PSNs are particularly suitable for targeting new analgesic treatments, especially at the levels of associated pathological DRG (54, 61). A recent study reports that central nervous system gene therapy by intravenous high-dose AAV causes asymptomatic and self-limited DRG inflammation and mild PSN degeneration in primates (62). Since these changes are very minor in comparison to the those induced by painful and neuropathic conditions that AAV injection would treat, this is unlikely to become a barrier to preclinical application of our approach.

In preclinical models, direct DRG delivery of AAVs encoding analgesic biologics can provide relief in chronic pain, with high transduction efficiency, flexibility for selective segmental localization, and minimal behavior changes attributable to the surgical procedure (63). In parallel, injection techniques are being advanced to achieve minimal invasive delivery of biologics for future clinical pain therapy (64, 65). Small peptides derived from the target protein sequences can serve as decoy molecules to selectively interfere with the function of their target signaling proteins by preemptively binding to them (66). We have successfully employed this strategy in rat models to induce analgesia by block of T-type/Ca_V_3.2 channel functions (13) and by blocking membrane trafficking of Ca_V_2.2 channels via interrupting its interactions with the structural protein of collapsin response mediator protein 2 (CRMP2) (66). Here, we extend the applicability of DRG-AAV strategy to the analgesic effectiveness of multiple PSN TTXs Na_VS_ blockade for neuropathic pain. These encouraging results that indicate efficacy and tolerability, if further validated for long-term efficacy and minimal side-effects, suggest the transformational potential of the approach for developing addiction free peripheral pain therapeutic agents. Beyond peripheral nerve injury-induced pain, dysfunctional Na_V_s have been found in various pain conditions, such as osteoarthritis (OA) that is frequently highlighted as an unmet medical need. Thus, for pain conditions like OA, targeting the TTXs Na_V_s might be therapeutically useful (67).

While our studies illustrate the power of rational analgesic peptide drug design strategy and provide encouraging results, we acknowledge several limitations in the current study. Different sodium channels traffic to distinct subcellular locations of PSNs (membrane, terminals, nodes of Ranvier, etc.), and the regulation of this process may provide a number of options to control neuronal excitability in different pathophysiological contexts. Injury-induced peripheral hypersensitization associated with Na_V_ malfunction affects multiple sites of the peripheral sensory nervous system, including augmented pain perception in the peripheral terminals, enhanced nociceptive signal transduction in PSN soma and T-junction, and increased neurotransmission in the spinal dorsal horn. At this early stage, our studies did not investigate differential actions by block of TTXs Na_VS_ along the pathway of peripheral nociceptors, nor did the results rule out the possibility that block of TTXs Na_V_s reduces pain by inhibiting afferent hyperexcitable input (68), thus indirectly modulating spinal cord and brain antinociceptive control circuits. Another limitation is that the molecular mechanism(s) of NaviPA1 functioning remains not fully investigated. The potential signaling pathways that the NaviPA1 affected could be many, since Na_V_1.7 PPI molecule networks involve multiple pathways and Na_V_1.7 (and other TTXs Na_V_s) intracellular segments serve as essential interfaces for many regulatory signaling molecules, including protein-lipids interactions (33, 34). Alterations of these molecules following nerve injury are essential for ectopic PSN hyperactivity and pain. Future work will address these questions.

## Materials and methods

### Animals

Adult male and female Sprague Dawley (SD) rats weighing 100-125g body weight (Charles River Laboratories, Wilmington, MA) were used. The estimated numbers of animals needed were derived from our previous experience with similar experiments (13) and a power analysis was not performed. The numbers of rats used were detailed in the relevant sections or figure legends of the experiments.

### Computational (*in silico*) designs

Rat full-length Na_V_1.7 aa sequence was retrieved from the UniProt KB knowledge database (UniProt Knowledgebase release 2018_11). Na_V_1.7 protein TM domains and intracellular termini and loops were predicted by Phobius (https://www.ebi.ac.uk/Tools/pfa/phobius/) (69). The Na_V_1.7 protein IDRs were predicted by analyzing the full-length Na_V_1.7 sequence using DEPICTER (DisorderEd PredictIon CenTER, http://biomine.cs.vcu.edu/servers/DEPICTER/) (17). Potential phosphorylation sites in the Na_V_1.7 full aa sequence were identified using Disorder Enhanced Phosphorylation Predictor (DEPP, http://www.pondr.com/cgi-bin/depp.cgi) (18). Potentially functional peptides within the IDRs were further analyzed SLiMPrints (http://bioware.ucd.ie/slimprints.html), which predict short linear motifs (SLiMs) based on strongly conserved SLiMs within IDRs (21). Peptide structure determination was analyzed by I-TASSER (https://zhanglab.ccmb.med.umich.edu/I-TASSER/) (23). MacVector ClustaIW (MacVector, Apex, NC) was used for vector designs and sequence alignments.

### Molecular cloning and AAV constructs

The construct pAAV-CBA-GFP-1.7iPAs encode the GFP-Na_V_1.7iPA fusion protein downstream a chimeric intron for enhancing transcription, driven by a hybrid CMV enhancer/chicken β-actin (CBA) promoter, and a mRNA stabilizing Woodchuck Posttranscriptional Regulatory Element (WPRE) sequence was inserted downstream of stop code of GFP-Nav1.7iPAs and upstream of human growth hormone poly A signals. Plasmids were subsequently used in transfection experiments and in AAV vector generation. To package AAV6-GFP-1.7iPA1, AAV6-GFPlinker, and AAV2/6-GFP-NP (a Na_V_1.7 N-terminal inert peptide) (referred to as AAV6-NaviPA1, AAV6-GFP, and AAV6-NP, respectively) for *in vivo* injection. AAV vectors were produced and purified in our laboratory by previously established methods (70). The titers (GC/mL) of AAV6-GFP), AAV6-NaviPA1, and AAV6-NP vectors were 2.45 x10^13^, 3.05 x10^13^, 2.64 x10^13^, respectively. Same batches of AAVs were used in all in vivo experiments.

### Cell culture

#### Cell lines

HEK293 cell line stably expressing human wide-type Na_V_1.7 (HEK1.7) was provided by Dr. Theodore Cummins. HEK293 cell lines stably expressing human wide-type Na_V_1.6 (HEK1.6), Na_V_1.3 (HEK1.3), Na_V_1.5 (HEK1.5), and CHO cells stable expression of human Na_V_1.8 (CHO1.8) were obtained from Charles River, Neuronal NG108-15 (NG105) neuronal-like cells and F11 cells (hybrid cells of mouse neuroblastoma cells with embryonic rat DRG neurons) were purchased from ATCC (Manassas, VA). These cells were using standard techniques, as described previously (13).

#### Generation of human Na_V_1.8 stable HEK293 cells

To generate Na_V_1.8 stable expression HEK293 cells (HEK1.8 cells), an expression plasmid of pcDNA3.1(+)-SCN10A-Furin-P2A-SCN2B was constructed (Genscript) in which a CMV promoter transcribes human Na_V_1.8α and Na²2 from a single open reading frame (ORF) expressing human *SCN10A* (NM_001293306.2) and *SCN2B* (NM_004588.5) separately linked by a 2A self-processing sequence derived from porcine teschovirus-1 (P2A) and a furin cleavage site (**Suppl**. **Fig**. **2**.) (71). Final construct was transfected in HEK293 cells, and the cells were frequently selected with G-418 (800 µg/mL), followed by establishment of single cell colonies using BIOCHIPS Single-cell Isolation Chip (ThermoFisher) according to the manufacturer recommended protocol. Na_V_1.8α and Nav²2 expression were determined by immunoblots using cell lysates, and prioritized by functional sodium current amplitude using conventional whole-cell voltage-clamp (see further).

#### Dissociated DRG neuronal culture

Dissociated DRG neuronal culture for EP was performed, as described previously (72) and were studied in 6∼8 h after harvest in EP experiments.

#### Human induced pluripotent stem cells (iPSC)-derived sensory neurons (hiPSC-SNs)

HiPSC-SNs, which were derived from female human ectoderm-neural crest stem cells, and ChronoTM Senso-MM complete growth medium were purchased from Anatomic. HiPSC-SNs maturing differentiation culture was performed per manufacturers’ recommendation.

#### Generation of lentivector expressing NaviPA1 and NP for hiPSC-SNs transduction

Lentiviral (LV) expression plasmid pWPT-GFP (73) was used to express NaviPA1 and NP (control) (**Suppl. Fig. 4**). LV were packaged using pWPT-GFPNaviPA1 and pWPT-GFPNP with packaging plasmid pCMVdR8.74 and envelop plasmid pVSV-g, concentrated, and products titrated in the range of 1×10^8^ to 2×10^8^ transduction unit/mL. Cultured hiPSC-SNs were infected by LV-LV-GFP, GFPNaviPA1 or LV-GFPNP in the presence of 8μg of polybrene (Sigma-Aldrich) per mL at an optimized multiplicity of infection:10.

### Electrophysiology (EP)

EP recordings were performed, as we described previously with minor modifications at room temperature (22∼25°C) (72, 74), in a blind manner where the electrophysiologist was not aware of the treatment. Patch pipettes, ranging from 1-2MΩ resistance, were formed from borosilicate glass (King Precision Glass Co., Claremont, CA) and fire polished. Recordings were made with an Axopatch 700B amplifier (Molecular Devices, Downingtown, PA). Signals were filtered at 5 kHz and sampled at 10 kHz with a Digidata 1440A digitizer and pClamp10 software (Molecular Devices, San Jose, CA). Series resistance (3–5MΩ) was monitored before and after the recordings, and data were discarded if the resistance changed by 20%. After achieving the whole-cell recording, capacitance (*C*m) and series resistance (*R*s) were compensated accordingly.

#### Sodium channel current (I_Na_) recording in cultured cell lines

Whole-cell voltage-clamp to recording **I_Na_** was performed in HEK1.7, HEK1.3, HEK1.6, HEK1.5, HEK1.8, CHO1.8, NG108-15 cells, and F11cells in current-density (I-V) and fast-inactivation voltage protocols. External solution consists of the following (in mM): 110 NaCl, 20 tetraethylammonium-Cl, 0.01 CaCl_2_, 0.1 CoCl_2_, 5 MgCl_2_, 10 HEPES and 5.56 mM glucose (pH 7.4, 310–315 mosM/L). The internal pipette solution consisted of (in mM): 10 NaCl, 130 CsCl, 5 MgCl_2_, 5 EGTA, 2.5 Na^2+^ATP and 10 HEPES (pH 7.2). After formation of a tight seal (maximal leak amplitude <150 pA), membrane resistance and capacitance were determined. The voltage dependence of activation was assessed from holding potential using 50 ms pulses (test-pulse) to a range of test potentials from −100 mV to +50 mV in 5 mV or 10 mV incremental steps with an interval of 5 s. Current density was calculated by normalizing maximal peak currents with cell capacitance. The voltage dependence of steady-state fast inactivation was measured using a two-step protocol. A 500 ms pre-pulse with various potentials ranging from *V*_hold_ to +10 mV in 5 or 10 mV incremental steps was used to inactivate the channels. This pre-pulse was immediately followed by a 40 ms test-pulse at 0 mV to determine the remaining fraction of available channels. Inward current measured during the test-pulse to 0 mV was normalized to the cell’s maximum test-pulse inward current. To determine the conductance-voltage (I–V) relationships of voltage-dependent activation, the peak current densities during each voltage command step were fitted to a smooth curve with a Boltzmann equation: I=G_max_(V-E_rev_)/[(1+exp[(V-V_50_)/k))], which provided the maximum conductance (G_max_). Normalized activation curves were fitted with a Boltzmann equation G/G_max_=1/(1+exp(V_50_-Vm)/k), where G was calculated as follows: G=I/(Vm-Erev). The steady-state inactivation curves were fitted with I/Imax=1/(1+exp-(V_50_-Vm)/k). In all the equations, V_50_ denotes the half-activation and half inactivation potentials, Vm is the membrane potential, Erev is the reversal potential, k is the slope factor, G is the conductance, and I is the current at a given Vm; G_max_ and I_max_ are the maximum conductance and current, respectively. Current density was obtained by dividing the maximum peak current (pA) by the cell capacitance (pF). Voltage errors defined by *Rs* x *Imax* were minimized by using 75∼80% series resistance compensation.

#### TTXs and TTXr I_Na_ recording in DRG dissociated neurons (male rats) and hiPSC-SNs

Isolated I_Na_ was recorded from single small/medium DRG neurons (¢.¢35 μm in diameter, 4wk after AAV-DRG injection in naïve rats) and differentiated hiPSC-SNs in bath solution that contained the following (in mM): 80 NaCl, 50 choline-Cl, 30 TEA-Cl, 2 CaCl2, 0.2 CdCl2, 10 HEPES, and 5 glucose, pH 7.3 with NaOH. internal solution containing the following (in mM): 70 CsCl, 30 NaCl, 30 TEA-Cl, 10 EGTA, 1 CaCl2, 2 MgCl2, 2 Na2ATP, 0.05 GTP, 10 HEPES, and 5 glucose, pH 7.3 with CsOH. A voltage protocol was adopted to separate TTXr I_Na_ and TTXs I_Na_ (26, 27). In brief, A 500 ms prepulse to **-**120 or **-**50 mV was applied before a 50 ms test pulse from **-**100 to 40 mV with steps of 5 or 10 mV by test pulses from **-**50 to 0 mV. Both TTXs and TTXr I_Na_ were apparent after the **-**120 mV prepulse; only TTX-R I_Na_ was obtained after the **-**50 mV prepulse, and the TTXs component was obtained by digitally subtracting the TTXr I_Na_ from the total I_Na_.

#### High-voltage activated (HVA) I_Ca_ and Voltage-gated potassium channel current (I_Kv_)

To record HVA I_Ca_ was recorded in rat DRG dissociated neurons (sham-operated, AAV6-GFP, AAV6-NP, and AAV6-NaviPA1 transduced neurons, 4wk after AAV-DRG injection), and I_Kv_ recording were conducted in non-differentiated NG108-15 cells, as described previously (13).

#### Whole-cell current-clamp recording on dissociated DRG neurons (male rats)

Whole-cell current-clamp recording of dissociated DRG neurons was performed, as described previously (22, 42, 72). Dissociated small- and medium-sized DRG neurons (¢.¢40μm in diameter) from sham-operated animals, rats with TNI only, and dissociated DRG neurons with clear GFP expression from TNI rats injected with AAV6-GFPNP or AAV6-NaviPA1 at 8-week after TNI and 6-week after vector injection were used for recording (n=5 rats per group). The membrane input resistance was calculated by dividing the ending amplitude of steady-state hyperpolarizing voltage deflection by the injected current (75). APs were generated by injection of a series of current pulses (180 to 280 pA in steps of 20 pA, 250 ms). The baseline potentials were recorded for 20 ms before the stimulus pulses were injected into the neurons. Resting membrane potential (RMP) was defined as the mean value of the 20 ms pre-stimulus potential in the first trial and the AP rheobase as the minimum current required to evoke the first AP. The neurons with stable resting membrane potentials (RMP) more negative than −40 mV and overshooting APs (**>**80 mV RMP to peak) were used for additional data collection. AP frequency was determined by quantifying the number of APs elicited in response to depolarizing current injections (250 ms).

### Microinjection of AAV vectors into DRG

AAV vector solution was microinjected into right L4 and L5 DRG using previously described techniques (13, 60). Rats received L4 and L5 DRG injections of either AAV6-NaviPA1 or AAV6-NP (one vector per rat), consisting of 2 μL with adjusted titers containing a total of 2.0 x10^10^ genome viral particles.

### Animal pain model and behavior testing

#### TNI

Animals were anesthetized using isoflurane at 4% for induction and 2% for maintenance. TNI surgery was performed as we described previously (13). Sham-operated rats were subjected to all preceding procedures without nerve ligation and transection.

#### Stimulated behavior testing

Behavioral tests were conducted between 9:00 AM and 12:00 AM, as we described previously (13). The experimenters were blinded to the treatment during all data acquisition. Stimuli were applied to the lateral margin of the plantar aspect of the foot in the sural area of innervation. Sensory tests included eliciting reflexive behaviors induced by von Frey test, Pin test, cold stimulation (acetone), and heat stimulation (Hargreaves test), and were carried out as previously described (60).

#### Gabapentin injection

Gabapentin (GBP, Sigma-Aldrich) was dissolved in saline immediately before injections and administered intraperitoneally (i.p.) at a volume of 0.5-1.0 ml (final dose=100 mg/kg body weight).

#### Conditioned place preference (CPP)

A 3-chamber CPP apparatus was used (Med Associates, St. Albans, VT) in which 2 sliding doors separate the central chamber from the 2 side chambers that have distinct wall stripes and flooring. The CPP procedure: 1) On day 1, rats were acclimated to the CPP boxes for 30 min, with open access to each of the three chambers. On the preconditioning day, rats were placed in the grey middle chamber and allowed to explore both sides of chambers for 15 minutes and the time spent in each side was recorded, and the preferred and nonpreferred chambers were identified. 2) On the conditioning days, place conditioning was conducted using a biased assignment approach to drug pairing: saline was paired with the preferred chamber in the morning, and GBP was paired with the non-preferred chamber in the afternoon with a 6 hr interval (injections were never paired with the middle grey chamber). Conditioning consisted of the following sequential steps: intraperitoneal injection and restriction of the animal within the preferred chamber (saline) or non-preferred chamber (GBP) for 45 min. We used a 45 min conditioning time based on tests that gabapentin maximally reduced mechanical hypersensitivity at 30–60 min after i.p. injection (13). Animals were conditioned for 2 days since 2-day GBP has been reported sufficient to produce CPP in rodent pain models (76, 77); 3) For postconditioning testing, the animals were placed back into the middle grey chamber of the CPP chambers with free access to all chambers for 15 minutes. The difference score for each animal was calculated, by subtracting the time spent in the saline-paired or GBP-paired chamber before pairing (during preconditioning) from the time spent in each chamber after pairing (postconditioning), and then averaged within each group. Each rat had only a single CPP test six weeks after AAV injection. A CPP is defined if the animals spend significantly more time in the GBP-paired chamber versus the saline-paired compartment.

### Immunofluorescent staining

The previously described protocol was adopted (73). Primary antibodies: mouse GFP (1:500, Santa Cruz Biotechnology, SCB, CA. sc9996), rabbit GFP (1:500, Cell signaling, Danvers, MA. 2555), rabbit Na_V_1.7 (1:400, Alomone, ASC008), rabbit Na_V_1.6 (1:400, Alomone, ASC009), rabbit glial fibrillary acidic protein (GFAP, 1:1000, Dako, CA, Z0334), goat myelin basic protein (MBP, 1:500, SCB, sc13912), mouse neurofilament (NF200, 1:1000, Sigma-Aldrich, N6389), and mouse ²3Tubulin (Tubb3, 1:500, SCB, sc-80016). The fluorophore-conjugated (Alexa 488 or Alexa 594, 1:2000) secondary antibodies (Jackson ImmunoResearch, West Grove, PA) were used to reveal immune complexes. The immunostaining was examined, and images were captured using a Nikon TE2000-S fluorescence microscope (El Segundo, CA) with filters suitable for selectively detecting the green and red fluorescence using a QuantiFire digital camera (Optronics, Ontario, NY). For measurement and quantification of immunostaining, positive marker antibody immunostainings were defined as the cells with the fluorescence intensity greater than average background fluorescence plus 2 standard deviations of the cells in an adjacent area in the same IHC slide of negative control (the first antibody omitted) under identical acquisition parameters (n=10 for different markers). NIH ImageJ software (http://rsbweb.nih.gov/ij/) was used for analysis.

### Immunoblot

Immunoblots were performed as described previously (13). To examine the subcellular localization of Na_V_1.7 in HEK1.7 cells, the cells were homogenized and then fractionated to obtain plasma membrane and cytosolic fractions using the ProteoExtract Subcellular Proteome Extraction Kit (Millipore, Billerica, MA). Antibodies: mouse GFP (1:1000), rabbit Na_V_1.7α (1:1000), rabbit Na_V_1.8α (1:1000, Alomone, ASC-016), rabbit Nav²2 (1:1000, Alomone, ASC-007), mouse Na^+^/K^+^ ATPase 1α (NKA1α, 1:600, SCB, sc514614), and mouse Gapdh (1:5000, Sigma-Aldrich, SAB1403850). Immunoreactive proteins were detected by Pierce enhanced chemiluminescence (ThermoFisher) on a ChemiDoc Imaging system (Bio-Rad) after incubation for 1 hr with HRP-conjugated second antibodies (1:5000, Bio-Rad).

### GFPNaviPA1 affinity pull-down followed by immunoblots, silver staining, mass spectrometry, and PIP strip assay

#### GFP affinity pull-down

This was performed using ChromoTek GFP-Trap kit. Briefly, HEK1.7 cells were transiently transfected to express GFPNaviPA1 or GFPlinker and/or GFP1.7iPA2 as controls. After 48 h, cells were lysed with ice-cold nondenaturing lysis buffer containing (in mM): 10 Tris/Cl pH 7.5, 150 NaCl, 0.5 EDTA, with 0.5 % Nonidet P40 substitute (a non-ionic and non-denaturing detergent), and protease inhibitor cocktail. The extracted cell lysates were diluted with 300 mL of dilution buffer containing (in mM) 10 Tris/Cl pH 7.5, 150 NaCl, 0.5 EDTA. For GFP-affinity pulldown, GFP-Trap Agarose Beads were equilibrated by adding 25 mL to 1.5 mL reaction tube with 500 mL of dilution buffer, cell lysates (2.5 mg total protein per sample) were incubated with GFP-Trap agarose by tube rotated end-over-end overnight at 4°C, followed by precipitation of beads by centrifugation at 12,000rpm.

#### Silver stain, Nav1.7 immunoblot, and mass spectrometry

The extracted cell lysates (input) and GFP-affinity pulldown beads (pulldown) were performed for 4-20% SDS-PAGE gel, followed by immunoblots using Na_V_1.7 antibody and silver stain on an additional SDS-PAGE gel using Pierce silver stain kit (ThermoFisher). The stained gel regions of interests were excised, and in-gel trypsin digested, as described previously (78). Extracted tryptic peptides were analyzed by nano reversed-phase liquid chromatography tandem mass spectrometry (nLC-MS/MS) using a nanoACQUITY (Waters Corporation, Milford, MA) online coupled with an Orbitrap Velos Pro hybrid ion trap mass spectrometer (ThermoFisher). The precursor ions were selected automatically by the instrument. Resultant MS/MS data were analyzed using the Mascot search engine (Matrix Science version 2.4) against the SWISSPROT human database.

#### PIP strip assay

NG108-15 cells transfected with GFPNaviPA1 and GFPlinker were prepared for cell lysate using a denaturing RIPA buffer containing (in mM): 10 Tris-HCl pH 7.5, 150 NaCl, 0.5 EDTA, with 0.1% SDS, 1% Triton X100, 1% deoxycholate, and protease inhibitor cocktail. The GFP-affinity pulldown beads from NG108-15 cells were size separated using 4-20% SDS-PAGE gels followed by silver stain. PIP strip assay (Echelon Biosciences, Salt Lake City, UT) was performed to analyze interactions between purified GFPNaviPA1 from NG108-15 cells and membrane lipids, compared to purified GFP from NG108 as control. In brief, PIP (phosphatidylinositol phosphate) strip membranes were blocked in 3% (w/v) fatty acid-free BSA (Sigma-Aldrich) in TBST (50 mM Tris-HCl, pH 7.5, 150 mM NaCl, and 0.1% Tween 20) for 1 h.

The membranes were then incubated in the same solution with the purified GFPNaviPA1 or GFP (equal 1.5 μg/mL) overnight at 4°C with gentle agitation. The membranes were washed 3 times over 30 min in fatty acid-free BSA-TBST. The membranes were incubated for 1h with 1:2000 dilution of HRP conjugated anti-GFP monoclonal antibody (Proteintech, HRP66002) at room temperature, then washed 6 times over 1 h in TBST, and the protein that was bound to the membrane by interaction with phospholipids was detected by enhanced chemiluminescence on a ChemiDoc Imaging system.

### Statistics

Statistical analysis was performed with GraphPad PRISM 9 (GraphPad Software, San Diego, CA). The methods were detailed in the figure legends and results were reported as mean and standard deviation of mean (SEM). Differences were significant for values at *p*<0.05. For comparisons between groups, in the pilot *in vivo* testing of TNI operation at three weeks after AAV intraganglionic injection, the effects of vector injection were characterized by treatment area under the curve (tAUC) analysis; in the treatment protocol of established pain, the measures immediately before AAV injection at the 14^th^ day post TNI were used as the tBL for calculating tAUC. In treatment of established male TNI pain, the data points in one rat who died on the second day after treatment AAV injection (no diagnostic report and likely due to surgical injury) were excluded from analysis.

### Study approval

All animal experiments were performed with the approval of the Medical College of Wisconsin Institutional Animal Care and Use Committee in accordance with the National Institutes of Health Guidelines for the Care and Use of Laboratory Animals.

## Supporting information

file:///Users/hongweiyu/Desktop/Catalyst/MS/MS1/Submit/JCI/Supplemental%20materials1.html

## Funding

This research was supported by a grant from a 2022 award from Dr. Ralph and Marian Falk Medical Research Trust, Bank of America, Private Bank (to HY and QHH); National Institutes of Health grant R61NS116203 (to HY and QHH); MCW Therapeutic Accelerator Program (2022) (to HY); and 2023 Advancing a Healthier Wisconsin Endowment Project (5520680) (to HY).

## Author contributions

HY designed the study and wrote the manuscript. HY, QHH, and TRC revised manuscript. SMS, BIZ, and HY performed most experiments, analyzed data, and organized all figures. QHH supervised DRG injection. CSQ, MR, UG, and FF participate and assist for experiments. HY and QH. obtained fundings. TRC provided HEK1.7 stable cell line and consulted EP experiments. All authors reviewed the manuscript.

## Data and materials availability

All experimental data generated or analyzed in this study are either included in this article (and its supplementary information files) or will be made available upon reasonable request.

## Notes

### Competing Interest Statement

The authors have declared no competing interest.

### Summary of Updates

Continue first submission

