## Supplementary material for "Peripherally targeted analgesia via AAV-mediated sensory neuron-specific inhibition of multiple pronociceptive sodium channels in rat": file:///Users/hongweiyu/Desktop/Catalyst/MS/MS1/Submit/JCI/Supplemental%20materials1.html

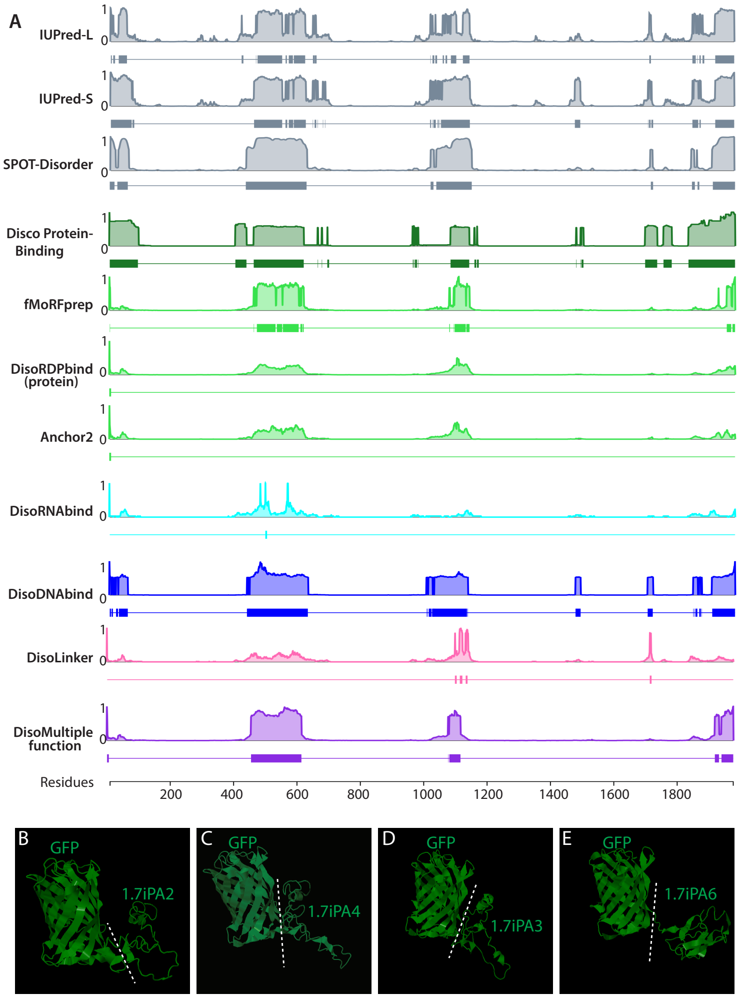


**Figure S1. In silico DEPICTER and I-TASSER**. (**A**) IDRs are predicted by DEPICTER [1], a prediction algorithm that aggregates results from the multiple servers (IUPred-L and -S: Prediction of Intrinsically Unstructured Proteins. SPOT-Disorder-Single: Accurate Single-Sequence Prediction of Protein Intrinsic Disorder. Disco Protein Binding: Prediction of IDR Protein Binding. fMoRFpred: Fast Molecular Recognition Feature predictor. DisoRDPbind: Predictor of disorder-mediated RNA, DNA, and protein binding regions. Anchor 2: Potential binding sites in disordered regions. DiscoRNA(DNA) Bind: Prediction of Potential RNA (DNA) binding regions. Disordered: Flexible Linker predictor. DiscoMultipleFunction: Annotations of disordered multifunctional residues), with each algorithm indicated on the left side. (**B-E**) The crystal structure analysis of GFP1.7iPA2, 3, 4, and 6 by I-TASSER, as indicated.

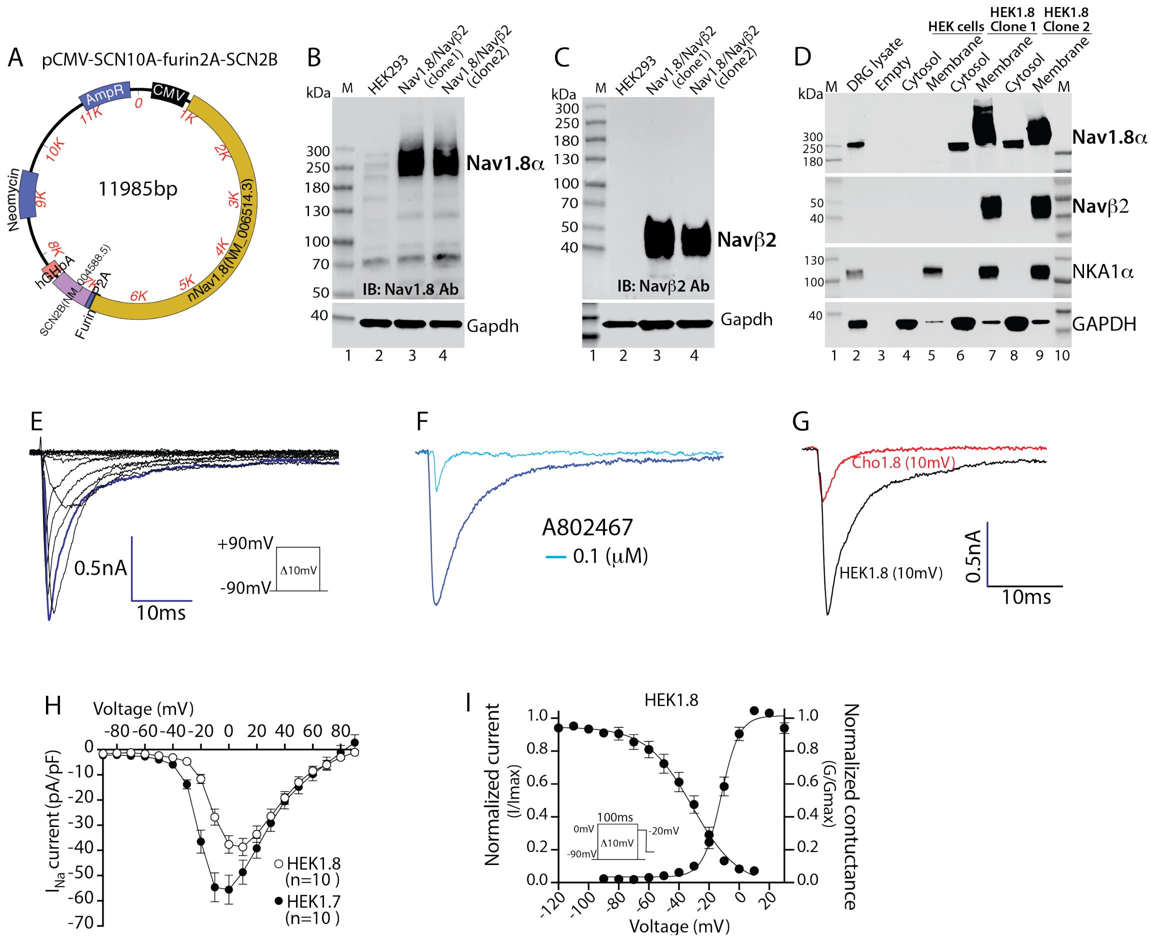


**Figure S2. Establishment of Nav1.8 stable expression system based on HEK cells. (A)** An expression plasmid of pcDNA3.1(+)-SCN10A-Furin-P2A-SCN2B was constructed. (**B**-**C**) Immunoblots of cell lysates verify stable expression of Nav1.8α (**B**) and Navβ2 (**C**), both are highly enriched in the plasm membrane (**D**). Traces of voltage-gated inward $I_{Na1.8}$ by whole-cell voltage-clamp recording (**E**), which show >85% inhibition by A802467 (100nM) (**F**) and comparison of single traces (+10mV) recorded from HEK1.8 cells and CHO cells stably expressing human Nav1.8 (**G**). (**H**) Comparison of I/V curves between HEK1.7 and HEK1.8 cells. (**I**) Voltage-dependent activation and steady-state inactivation curves recorded from HEK1.8 cells. Inserts are the current/time scale and recoding protocols.

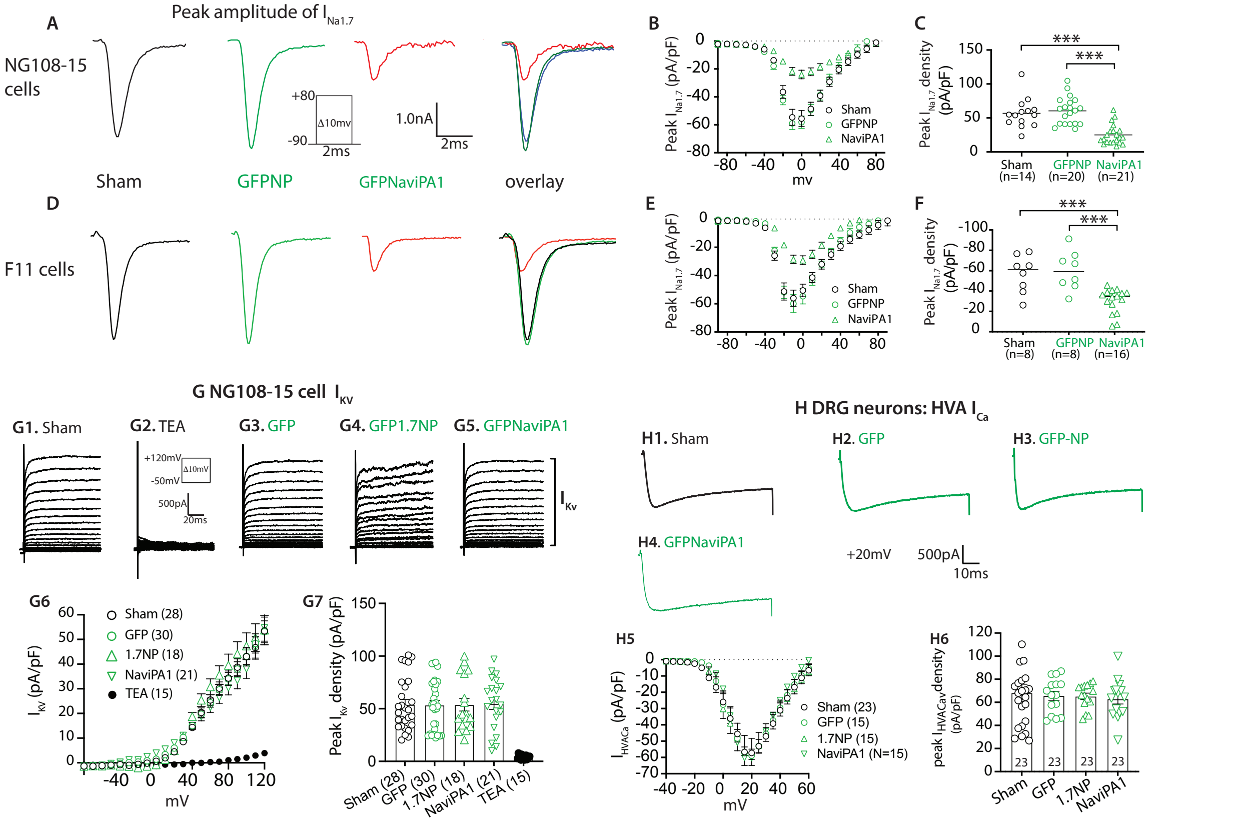


**Figure S3. NaViPA1 on $I_{Na1.7}$ of NG108 and F11 cells, $I_{Kv}$ of NG108 cells, and HVA $I_{Ca}$ of DRG neurons**. Representative $I_{Na1.7}$ single traces at 0 mV recorded from sham-, GFPNP-, NaViPA1-transfected cells, merged $I_{Na1.7,}$ I/V curves, and peak $I_{Na1.7}$ density from NG108-15 cell (**A-C**) and F11 cells (**D-F**). Insets: protocol and current/time scales. ***p<0.001, one-way ANOVA and turkey post hoc. (**G**) Representative $I_{Kv}$ of sham-NG108 cells showing $I_{Kv}$ defined by outward currents blocked by Tetraethylammonium (TEA, 5mM) or NG108 cells transfected with GFP, 1.7NP, and NaViPA1 (**G1-G5**). insets: recording protocol and current/time scales. $I_{Kv}$ density-voltage (I/V) curves (**G6**) and quantitative analysis of peak $I_{Kv}$ density (**G7**), $p>0.05$, one-way ANOVA and Tukey *post hoc.* (**H**) DRG neuron HVA $I_{Ca}$ recording. Typical HVA $I_{Ca}$ trace in a small-sized neuron from a naïve rat shows a threshold for activation around −30 mV and a maximum current amplitude activation at -10mV, displaying small inactivation (**H**). Typical traces of HVA $I_{Ca}$ recorded at -10mV of neurons from a sham-operated rat (**H1**) and naïve rats injected with AAV6-GFP (**H2**), -1.7NP (**H3**), and -NaViPA1 (**H4**). HVA $I_{Ca}$ density-voltage (I/V) curves (**H5**) and averaged peak HVA $I_{Ca}$ density (**H6**), $p>0.05$, one-way ANOVA and Tukey *post hoc*.

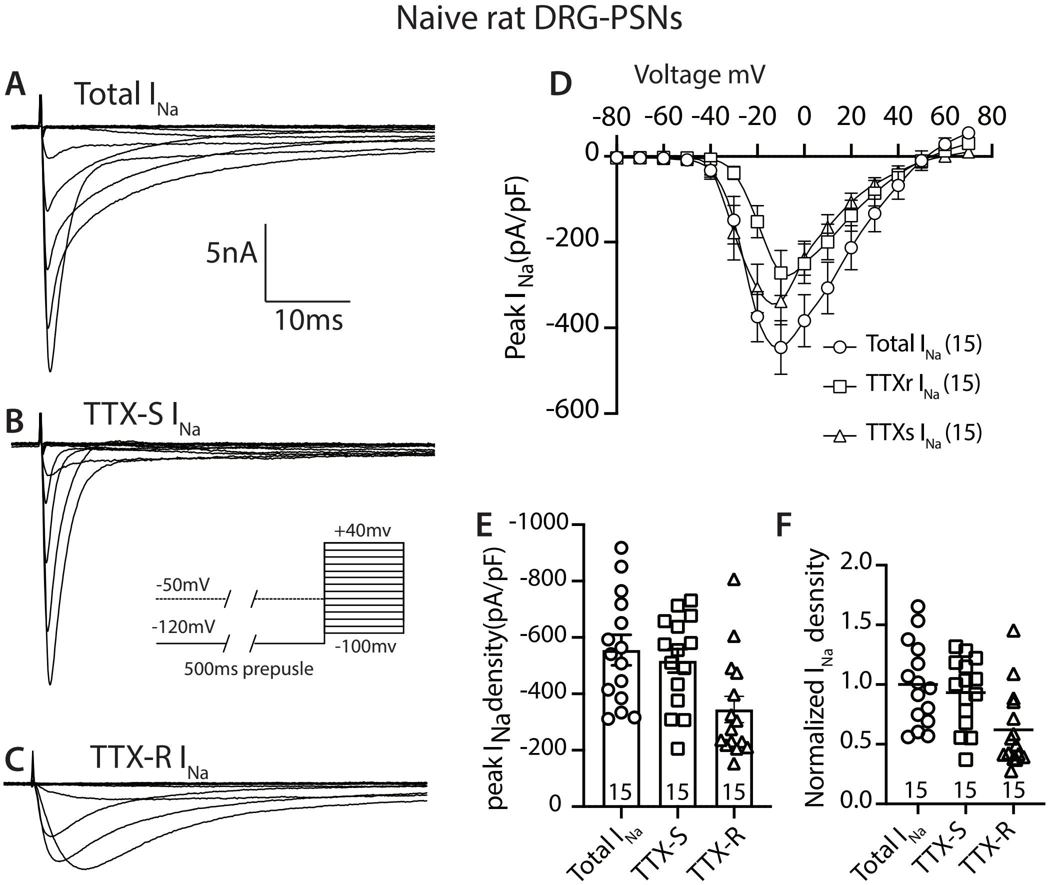


**Figure S4**. **TTXs and TTXr $I_{Na}$ recording in DRG neurons from naïve rats**. (**A-C**) Representative trances of voltage gated total, TTXs, and TTXr $I_{Na}$ recorded from small-sized DRG neurons. Protocol for separation of TTXr and TTXs $I_{Na}$: A 500 ms prepulse to -120 or -50 mV was applied before a 50ms test pulse from-100 to +40mV with steps of 10 mV(inset). Both TTXs and TTXr $I_{Na}$ were apparent after the -120 mV prepulse (top traces); only TTXr $I_{Na}$ were obtained after the -50 mV prepulse (bottom traces), and the TTXs component was obtained (middle traces) by digitally subtracting the TTXr $I_{Na}$ from the total $I_{Na}$. (**D**) Average peak $I_{Na}$ density–voltage relationships for total, TTXs, and TTXr $I_{Na}$ of small DRG neurons. Smooth lines are I–V curves generated using the Boltzmann fit parameters of the respective activation curves. Averaged peak $I_{Na}$ densities (**E**) and normalized peak $I_{Na}$ densities (**F**) of total, TTXs, and TTXr $I_{Na}$.

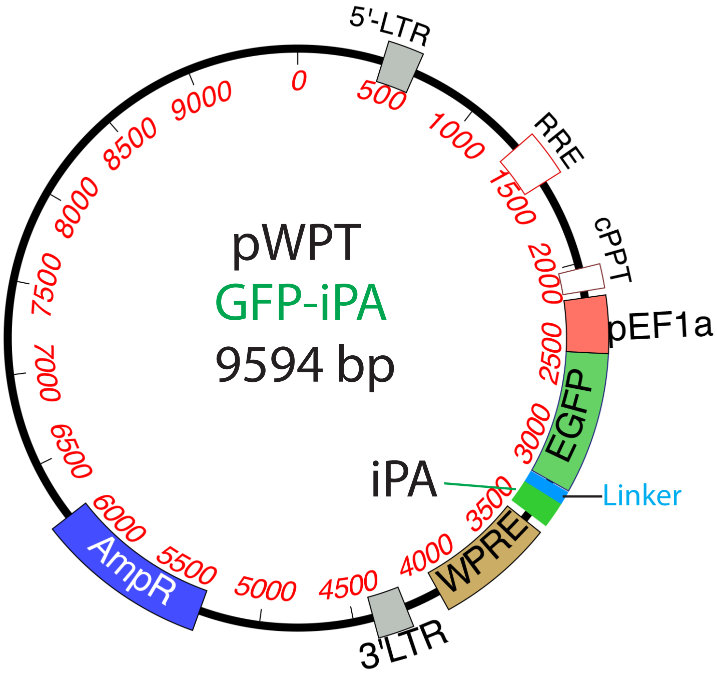


**Figure S5. Lentivector construct to express NaviPA and NP**

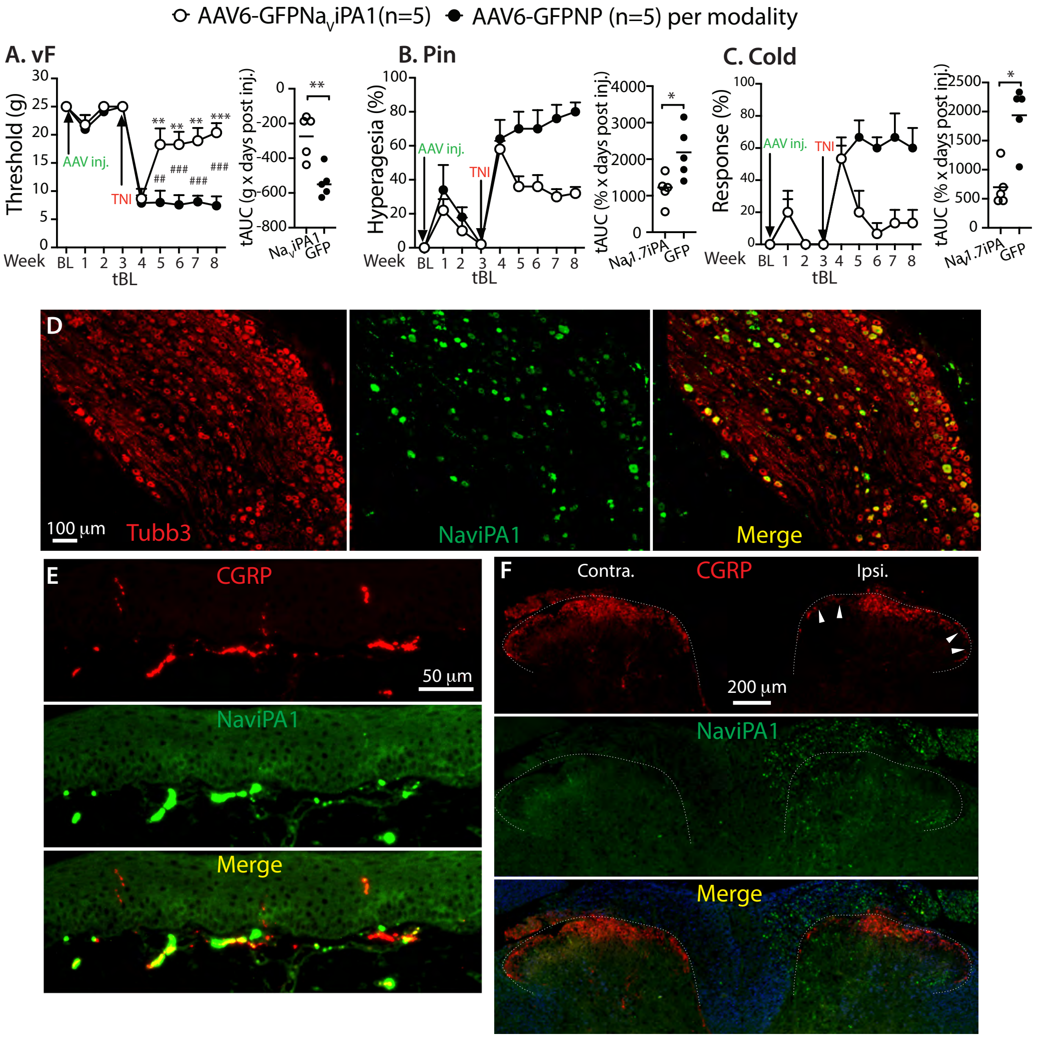


**Fig. S6. In vivo pilot analgesic testing of NaviPA1**. (**A-C**) Time courses for the group averages of sensitivity to vF, Pin, and Cold after DRG injection of either AAV6-NaviPA1 (n=5) or AAV6-NP (control, n=5) and subsequent TNI induction 3-wk after AAV injection as indicated; $^{**}p$<0.05 and $^{***}p$<0.001 for comparisons to the TNI BL (tBL) within group, and $^{\#\#}p$<0.01 and $^{\#\#\#}p$<0.001 for comparisons between groups. Repeated measures parametric two-way ANOVA for vF and Heat followed by Tukey (within group) and Bonferroni (between group) *post hoc*; and non-parametric Friedman ANOVA for Pin and Cold tests and Dunn's *post hoc*. Right panels of A-C show TNI AUC (tAUC) calculated using measures 21-day post AAV and before TNI as tBL; * $p$<0.05 and **$p$<0.01, comparisons of tAUC between groups (unpaired, two-tailed Student's $t$ tests for vF, and Mann–Whitney U tests for Pin and cold). Representative montage IHC images of DRG section (**D**, colabeled GFP with Tubb3, showing colocalization in merged image), ipsilateral (ipsi.) hindpaw skin section (**E**) and spinal cord section (**F**), colabeled GFP with CGRP (red), showing colocalization in merged image (arrowheads point to reduced CGRP innervation in ipsi. DH).
